# The DivisionCounter, a method for counting large ranges of cell divisions *in vivo,* reveals cell dynamics of leukemic cell killing via CAR-T therapy

**DOI:** 10.1101/2024.06.24.600543

**Authors:** Lucie S P Hustin, Cecile Conrad, Chang Liu, Jaime Fuentealba, Silvia Menegatti, Seva Shneer, Aude Battistella, Fanny Tabarin, Tom S Weber, Sebastian Amigorena, Ken R Duffy, Leïla Perié

## Abstract

Cell division drives multicellular growth and its dysregulation can cause disease. While approximately 44 divisions are needed to produce without death all 10^13^ cells in the human body, current methods are limited to count 10 cell divisions *in vivo* across diverse mammalian cell types. Here we introduce the DivisionCounter, a method to count cell division *in vivo* over large division ranges (∼70 divisions) using an easy fluorescence readout. We demonstrate that the DivisionCounter accurately measures the average cell division count of cells both *in vitro* and *in vivo*. Its use revealed that leukemia tumor division rates are independent of the organ’s specific microenvironment and CAR-T cell treatment, providing an estimate of tumor death rates *in vivo*. The DivisionCounter method holds unique potential for quantifying contributions of cell division, death, and migration to the growth of healthy and pathological mammalian tissues.

## Introduction

Cell proliferation drives the growth of multicellular organisms during embryogenesis, tissue homeostasis, and regeneration after damage, and its dysregulation can lead to disease such as cancer. It is well established that: i) cells are limited in their number of times they can divide, known as the Hayflick limit, of 40-60 divisions^1^; ii) cells accumulate genetic mutations during division, which can eventually lead to oncogenesis^2^; and iii) the memory of previous divisions regulates the decision of stem cells to differentiate^3^. In contrast, we still know little about how many cell divisions are required to produce and maintain each organ in the body, or how cell divisions are regulated mechanistically. This lack of knowledge about tissue development in healthy or pathological processes is largely due to a lack of high-resolution tools for counting cell divisions *in vivo*.

Currently available methods for quantifying the number of cell divisions in mammalian cells are limited in resolution and biological applications (Table 1). One significant issue is that, while cell counting is often used as a proxy for cell division, it is unable to deconvolve cell death from cell division and so it measures the net cell population doubling rather than quantifying the number of cell divisions^4^. Live microscopy of cells can trace at most a few divisions in mammalian cells *in vivo* or tens of divisions *in vitro*, relying on direct observation of divisions^5^. Division diluting dyes, such as CFSE and CTV, use the dilution of a dye by two at each division to infer division counts *in vitro* and *in vivo*^6^. However, these dyes are limited to count 10 divisions at best due to the geometric dilution of the dye with division. Other fluorescence dilution–based methods have a resolution of 5 or 6 divisions^7,8^. Lengthier inference of division number has been realized by quantifying telomere lengths^9^, but this method is restricted to a few specific cell types without telomerase activity^10^. There is, therefore, a need for additional methods that can quantify more than 10 cell divisions in a broad range of mammalian cell types.

**Table 1.** DivisionCounter and state of the art technologies to count cell divisions.

| Technology | Main use<br>(*In vitro*/<br>*in vivo*) | Division<br>count of ... | Maximum<br>Value<br>(division count) | Target eukaryotic<br>cell types or<br>organism | Selected References |
| --- | --- | --- | --- | --- | --- |
| Thymidine<br>analogue | both | Single cell | 1 | All | (Sanders et al., 2017) |
| Time-lapse<br>microscopy | *In vitro*<br>*In vivo* | Single cell | ~10<br>~5 | Growing *in vitro*<br>*C. elegans* and<br>zebrafish | (Hawkins et al., 2009; Loeffler et al., 2019; Plambeck et al., 2022)<br>(Giurumescu et al., 2012; Sulston et al., 1983) |
| Tet-Off-H2B-FP | both | Single cell | ~5 | GMO organisms | (Bernitz et al., 2016; Wilson et al., 2008) |
| Cell tracer dyes | both | Single cell | ~10 | Haematopoietic<br>cells (primarily<br>lymphocytes) | (Grinenko et al., 2018; Horton et al., 2018; Lyons, 2000; Lyons and Parish, 1994) |
| Telomere length | both | Cell<br>population | ~100 | Telomerase<br>negative cells | (Harley et al., 1990; Rufer et al., 1999; Vaziri et al., 1994) |
| DivisionCounter | both | Cell<br>population | 100+ | Prone to lentiviral<br>infection | This work |

Here we present a novel methodology, the DivisionCounter, that can count more than 70 divisions, using a simple and low-cost fluorescence readout, that builds on previous qualitative recorders of cell proliferation^11,12^. Briefly, the DivisionCounter couples changes in fluorescent protein expression, that are induced by microsatellite instability, with cell division to compute the division counts of a cell population, through the use of a genetic construct and a mathematical formula. We demonstrate that the DivisionCounter can count, in a robust, quantitative manner, up to 74 cell divisions in murine and human cell lines. Using a Nalm6 human B-cell acute lymphoblastic leukemia xenograft mouse model, we show that the DivisionCounter can be used to study cancer cell division *in vivo* during cancer development in different microenvironments, including under treatment with CAR-T cell therapy. We find that Nalm6 leukemic cells divide *in vivo* at a rate of 1 division/day in immunodeficient NOD/SCID/IL-2Rγ-null (NSG) mice. Further, these cells have similar *in vivo* average division counts in five distinct organs (bone marrow, liver, lungs, blood and spleen), demonstrating that their proliferation is independent of the organ microenvironment. With adoptive CAR-T cell therapy, Nalm6 cells continued to divide with the same *in vivo* division rate despite the decline in their absolute counts due to CAR-T cell killing, resulting in a war of attrition between the tumour and the T cells. Thus, the DivisionCounter method enables not only the quantification of division rate *in vivo* but also estimation of tumour death rate based on the average division counts and cell counts, enabling the study of tissue growth modulation.

## Results

### The DivisionCounter method includes a genetic construct and a mathematical formula

To develop a method that can count more than 10 cell divisions, we decided to compute the average division count of a cell population solely using a division-dependent change in the fluorescent protein (FP) expression of a cell. To this end, we engineered a genetic construct incorporating a microsatellite upstream of the start codon of an in-frame FP (Fig. 1a), which we termed the DivisionCounter construct. This construct leverages the properties of microsatellites to change nucleotide length during DNA replication^13^. Consequently, there is a small probability at each cell division that the FP will shift out of its reading frame (Fig. 1a), leading to loss of its expression. In turn, the FP can also return to its reading frame during subsequent divisions with further changes in microsatellite length. As a result, the DivisionCounter construct displays three distinct states: one with FP expression and two with no FP expression (Fig. 1a). Briefly, the DivisionCounter construct is integrated into the genome of cells using a lentivirus, and cells are then sorted to give only FP+ cells; this 100% FP+ population of cells then loses the FP expression as they divide. Evolution of the out-of-equilibrium system of three states results in the cell population converging to 33% FP+ cells, with the remaining cells in one of the two other states of the construct (Fig. 1b). The DivisionCounter method uses the percentage of FP+ cells as a readout, measured by fluorescent detection methods such as flow cytometry, to compute the division count via a new mathematical formula between the average division count, on one hand, and percentage of FP+ cells and the probability of losing the fluorescence, on the other.

**Figure 1:**
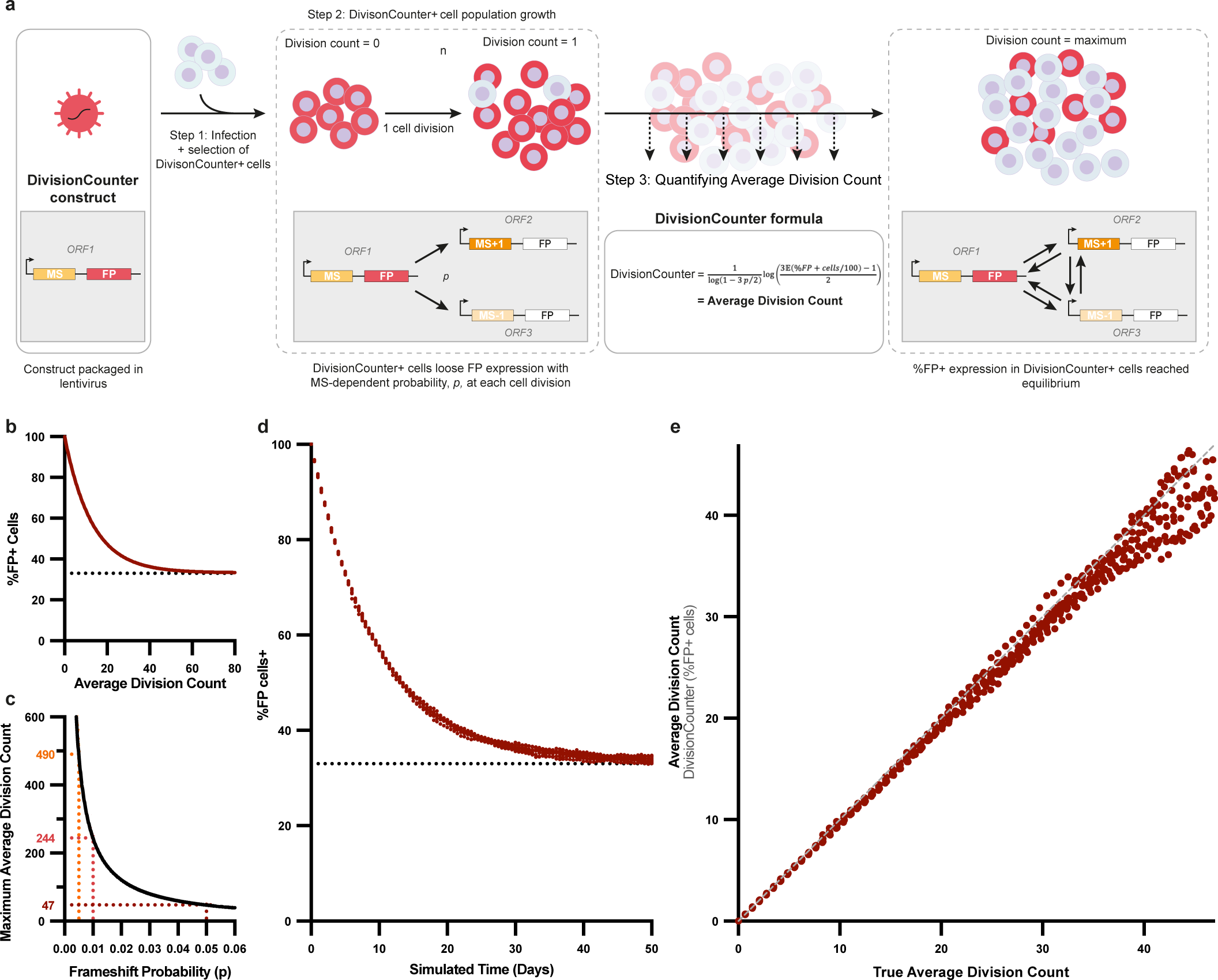
Concept of the DivisionCounter method. a) Schematic of the DivisionCounter method. First a DivisionCounter construct possess two essential features a microsatellite (MS) and a fluorescence protein (FP) in-frame, in open reading frame 1, (ORF1), all packaged in a lentivirus. After infection, the cells that are FP-positive are selected to start the DivisionCounter experiment. With each cell division, during DNA duplication, each cell has a chance, p, to undergo polymerase slippage on its MS sequence that will result in altering the MS length, pushing the FP out of frame (in ORF2 or ORF3). Reversibly, the FP in ORF2 or ORF3 can subsequently come back to its original reading frame (ORF1) during later cell divisions due to new slippage events. Finally, an equilibrium between the three states (ORF 1 to 3) of the construct’s FP is reached resulting in no more change of the percentage of FP+ cells. Therefore, FP+ DivisionCounter-infected cells lose their fluorescence with a probability, p, at each division until they reaches the equilibrium. The DivisionCounter method takes advantage of this cellular behavior to quantify the average division counts of DivisionCounter-infected cell population using the DivisionCounter formula at different time points only using p and the percentage of FP+ cells. b) DivisionCounter theoretical evolution of the percentage of FP+ cells over cell division count with a 0.05 FP loss probability. The percentage of FP+ cells decreases as cells divide to reach a plateau at 33% - the construct FP expression equilibrium point between its 3 ORF states. c) Maximum average division count for different probabilities of FP loss due to microsatellite slippage, computed using the DivisionCounter formula when the percentage PF+ cells reaches 35%. d) Results of the stochastic simulations of the loss of a FP+ with a probability of FP loss of 0.05 at each division over time. N= 100 simulations with parameters from Table S10. e) Average division count computed from the simulated % of FP+ cells against the true average division count in the simulations. N= 100 simulations.

To compute the division count of a cell population from the measurement of the percentage of FP+ cells, building on our previous theoretical work for irreversible FP changes^14–16^, we derived a novel formula that incorporates the three states into the calculation of the average number of divisions (Fig. 1a, see method). Assuming equal likelihood of transitioning between the three states during division, mathematical analysis predicts that the equilibrium state is for 1/3 of the cell population to be expressing the FP (Fig. 1b). Knowing the probability per division of losing the FP expression and a measurement of the percentage of FP+ cells, the newly derived DivisionCounter formula allows for the computation of the average division count of the cell population (Fig. 1b; see Extended Data Fig. S1a for different probabilities). The maximum division count determined by the DivisionCounter method is defined as the division count at which the FP expression border on the 1/3 equilibrium. For the probabilities of losing the FP expression of 0.05, 0.01 and 0.005, which are in the known range for microsatellite mutations^17–19^, the evaluation range of the DivisionCounter extends to 47, 244, and 490 divisions respectively (Fig. 1c), surpassing the 40 to 60 divisions of the Hayflick limit^1^. Of note, while the average division count of a cell cannot be quantitatively inferred after the system reaches equilibrium, the DivisionCounter provides a low boundary on the number of divisions made by the cell population.

Finally, we validated the DivisionCounter’s formula using stochastic simulations for different probability of FP expression loss (Fig. 1d-e and Extended Data Fig. S1b-c). In the simulated system, the percentage of FP+ cells decreased over time, as predicted by the DivisionCounter formula (Fig. 1d and Extended Data Fig. S1b). In addition, the DivisionCounter average division count obtained by inputting the percentage of FP+ cells in the DivisionCounter formula was highly correlated with the ground-truth average division count of these cells (Fig. 1e and Extended Data Fig. S1c). Thus, the DivisionCounter method has the capacity to accurately compute the average division counts of a cell population, surpassing the current *in vivo* limit of 10 divisions from the state-of-the-art methods.

### Measurement and inference of cell divisions *in vitro*

To assess the performance of the DivisionCounter method in a biological system, we infected HEK293T cells with either a lentivirus containing a 26-guanine (26G) microsatellite followed by the FP mScarlet (herein, 26G-DivisionCounter) or with a control lentivirus in which the microsatellite was replaced by a linker sequence of 26 nucleotides (26N-construct) (Fig. 2a). Using a low multiplicity of infection, we integrated one copy on average of the DivisionCounter construct per cell, resulting in approximately 10% of the cells being infected (Extended Data Fig. S2a, b). We obtained clear expression of mScarlet for both conditions (Extended Data Fig. S2b). After 3 to 7 days of culture, mScarlet-positive cells were sorted and grown in culture for 46 days (Fig. 2b). At each splitting, cells were counted, and a fraction of the cells was retained for a DivisionCounter readout, by recording the percentage of mScarlet-positive cells by flow cytometry (Fig. 2b). As a comparison control, cells were counted to compute cell population doublings, which is equivalent to the average division count in this culture condition for which there is no, or only minimal, cell death (Extended Data Fig. S2c). As predicted by the DivisionCounter method, the percentage of mScarlet-positive cells infected with the 26G-DivisionCounter decreased as the population proliferated, in contrast to cells with the 26N-construct (Fig. 2c). The changes in the percentage of mScarlet-positive cells followed the predicted pattern (Fig. 1b), decreasing with the average cell population doubling and reaching a plateau at 33% (Fig. 2c), corresponding to the three-state equilibrium of the DivisionCounter method. This behaviour was robust and reproducible across independent experiments (Extended Data Fig. S2d) and gave a value of the probability of fluorescence loss of 0.047 (CI= 0.044-0.049, Fig.2c). To further verify that the plateau at 1/3 corresponded to the three-state equilibrium of the DivisionCounter construct, HEK293T cells that had lost mScarlet expression were sorted, cultured, counted and assessed for their mScarlet expression for 39 days. The percentage of mScarlet-positive cells increased with cell divisions, until they finally reached a 1/3 equilibrium (Fig. 2c), experimentally confirming the appropriateness of the three-state hypothesis used in the DivisionCounter method.

**Figure 2:**
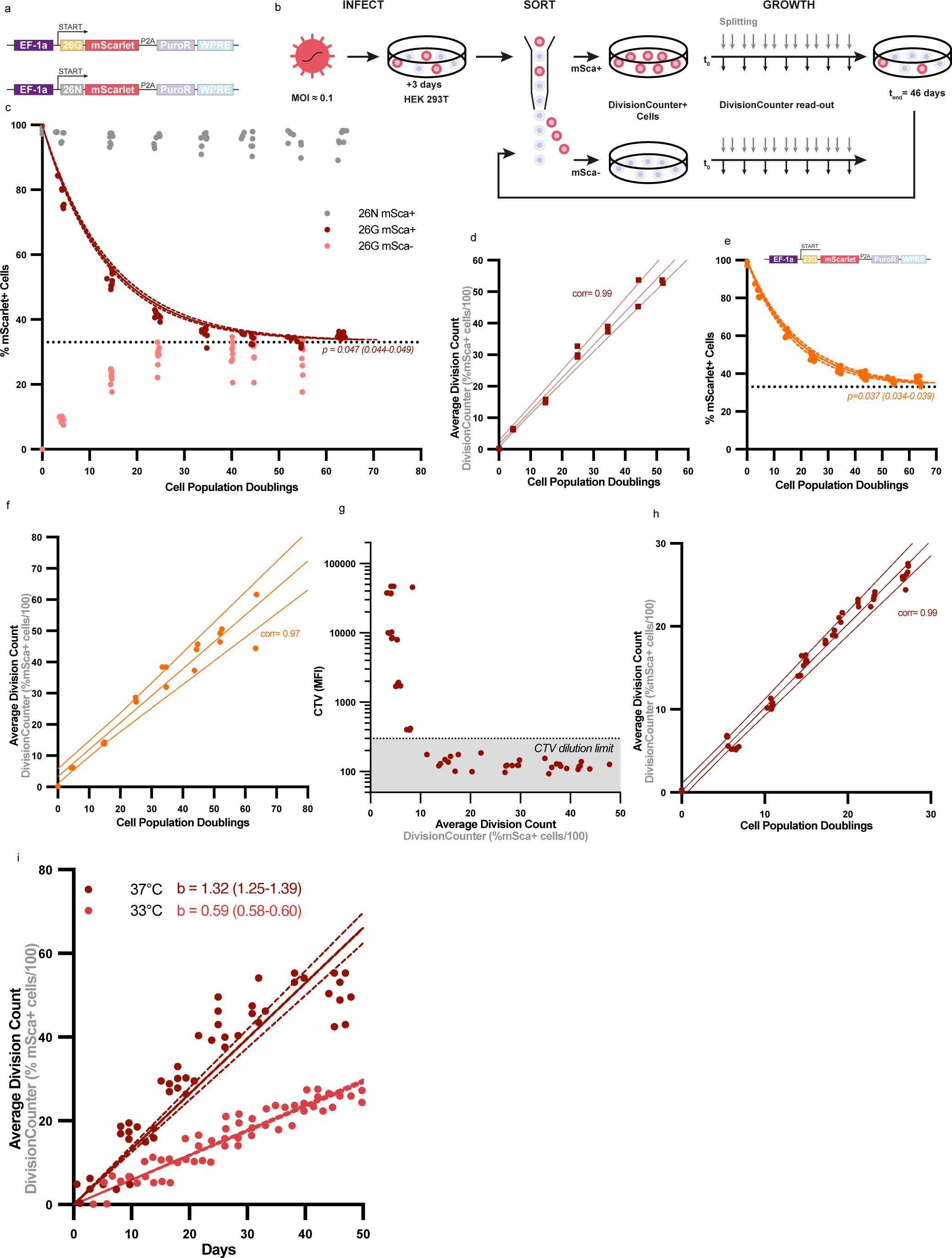
*in vitro* validation and benchmarking of the DivisionCounter method. a) 26G- and 26N-DivisionCounter constructs used in this figure. After the EF1-1a promoter and a start codon, a 26 guanines (26G) microsatellite or a linker sequence of 26 nucleotides as a control (26N) was placed in front of the fluorescent protein mScarlet, followed by the P2A self-cleaving peptide and Puromycin resistance gene (PuroR). b) HEK 293 T cells were infected with the 26G- or 26N-DivisionCounter lentivirus at a low multiplicity of infection. After 3 days, mScarlet+ (mSca+) cells were sorted and cultured for 46 days. Once or twice a week when cells were split, cells were counted and analysed for their mScarlet expression by flowcytometry as a read-out of the DivisionCounter method. Later, cells that became mScarlet-negative were also sorted, cultured and analysed for their mScarlet expression over time. c) Percentage of mScarlet-positive cells over the average division counts measured using cell counts (population doublings) for the 26G-DivisionCounter (red) and 26N-DivisionCounter (grey). For 26G-DivisionCounter, starting cell population were sorted to be either mScarlet-positive (dark red, mSca+) or mScarlet-negative (lighter red, mSca-). Each point represents an independent acquisition, from 3 experiments done in triplicates. The probability of losing mScarlet expression, p, was fitted by a nonlinear mixed-effects model using 3-fold cross-validation method. The full line shows the best fit with confidence interval obtained from bootstrapping. d) The average division count computed using the DivisionCounter versus the population doublings computed from cell counts for the 3^rd^ experiment. The percentage of mScarlet-positive cells are from the 3^rd^ experiment and the probability fitted in from the two other experiments as the 3-fold cross validation method. The Pearson correlation is reported on the graph (corr). Each point represents independent triplicate acquisition. e) same as in a and c but for the 23G-DivisionCounter. f) same as in d but for the 23G-DivisionCounter. g) Median Fluorescence Intensity of the Cell Trace Division in 26G-DivisionCounter infected HEK 293 T cells versus the DivisionCounter average division counts computed using the 26G-DivisionCounter. Grey area represents the limit of detection of CTV, defined by background expression of unstained HEK 293 T. Each point represents an independent acquisition from 2 experiments. h) 26G-DivisionCounter infected HEK 293 T cells were set to growth at 33°C. The average division count was calculated using the percentage of mScarlet-positive cells at 33°C into the DivisionCounter formula with the probability of mScarlet loss of the 37°C used in d versus the population doubling computed from cell counts at 33°C. The Pearson correlation is reported on the graph (corr). Each point represents independent acquisition from 3 experiments done in triplicate. i) DivisionCounter average division count change over time in HEK 293 T cells cultured at 37°C (dark red) and 33°C (light red). The probability of mScarlet loss used in DivisionCounter formula comes from inference made on HEK293T cells cultured at 37°C in (d). Each point represents independent acquisition from 3 experiments done in triplicate. The full lines represent the best fitted linear model and the dotted lines the confidence interval for the model. b= division rate obtained by linear regression and confidence interval.

To explore the accuracy of the DivisionCounter, we compared the average division count of DivisionCounter with the gold standard of cell population doublings (i.e., the average division obtained by cell counting). To this end, we used a 3-fold cross-validation method to fit the probability of losing mScarlet-expression using part of the data, which was then used to compute the average division count obtained with the DivisionCounter formula for the rest of the data. The obtained average division count was then compared with the cell population doublings. The probability of losing mScarlet-expression was fitted using a nonlinear-mixed-effects model to the loss in percentage of mScarlet-positive cells over the average cell population doubling (Extended Data Table S1). After repeating this cross-validation procedure several times, we observed a strong correlation between the average division counts obtained with the DivisionCounter and the average divisions obtained by cell counts (Fig. 2d and Extended Data Fig. S2e-f), validating that the DivisionCounter accurately counts divisions. Further, the 26G-DivisionCounter counted up to 56 divisions, with a confident interval (CI) of 54 to 58 divisions, thereby far exceeding the working range of other state-of-the-art methods.

We then explored the robustness, versatility and general applicability of the DivisionCounter by testing other microsatellite lengths and different cell lines. First, we infected cells with a DivisionCounter construct bearing a 23G microsatellite rather than a 26G microsatellite. Conducting the same experiment as before with infected HEK293T cells, we recapitulated all major behaviors of our DivisionCounter method: the percentage of mScarlet-positive cells decreased with the average cell population doubling and reached a plateau at 33% (Fig. 2e), validating the robustness of the DivisionCounter method. Performing the same bootstrapping procedure as for the 26G-DivisionCounter, we confirmed that the average division counts obtained with the 23G-DivisionCounter strongly correlated with the cell population doublings (Fig. 2f and Extended Data S2g-h). As the mutation rate of the guanine repeat–microsatellite increases with the number of guanine repeats^20^, we expected the 23G microsatellite to mutate at a lower rate than the 26G microsatellite, thereby increasing the maximum division number that can be counted. Indeed, we counted more divisions with the 23G- DivisionCounter than with the 26G-DivisionCounter, reaching a maximum of 74 divisions (CI = 69 to 79) as compared to 56, respectively (Fig. 2c, e), showcasing the versatility of the DivisionCounter method. That is, by changing the length of the microsatellite, the division range can be fine-tuned to suit the biological context. We next explored the DivisionCounter general applicability across different cell types. After infecting and sorting mouse embryonic fibroblasts (MEFs) and human triple-negative cancer cells (MDA-MB-468) with the 26G-DivisionCounter construct, we observed similar results to those seen in HEK293T cells (Extended Data Fig. S2i-j). The percentage of mScarlet-positive cells decreased with the average cell population doubling, consistent with the expected DivisionCounter behaviour. Notably, this decrease occurred more slowly than in HEK293T cells, enabling a higher number of divisions to be counted in these cell lines.

To further validate the DivisionCounter method, we compared its performance to another state-of-the-art division counting method: the Cell Trace Violet (CTV) dilution dye. MEF cells were double-labelled with the 26G-DivisionCounter construct and CTV then cultured, counted and assessed for their mScarlet and CTV status by flow cytometry over 51 days (Fig. 2g). Over the lower range of divisions that can be accurately followed with the two methods (i.e., below 10 divisions), we observed that the DivisionCounter average division count strongly correlated with that of the CTV dye dilution (Spearman coefficient = –0.73) (Fig. 2g), which further demonstrated the reliability and accuracy of the DivisionCounter method. Notably, however, CTV reached its dilution limit after 10 divisions, whereas the DivisionCounter counted up to 47 divisions (Fig. 2g), directly showing that the DivisionCounter method outperformed the dye dilution state-of-the-art method.

Finally, to confirm that the DivisionCounter mScarlet expression was linked to cell division, we modulated cell growth by culturing HEK293T cells infected with the 26G-DivisionCounter at 33°C to slow down cell division^21^. We first confirmed that cells cultured at 33°C grew slower than those at 37°C with no impact on cell mortality (Extended Data Fig. S2c,k) then assessed the percentage of mScarlet-positive cells over 49 days in *in vitro* culture. Using the DivisionCounter formula and the probability of mScarlet loss from HEK293T cultured at 37°C to compute the average division count, we predicted accurately the cell population doublings, as shown by their high correlation (Fig. 2h). These results further validated that changes in the percentage of mScarlet-positive cells were exclusively linked to cell division. Importantly, the DivisionCounter also allowed us to quantitatively measure the division rate. Fitting a linear regression to the average division counts over time, we estimated that cells cultured at 33°C grew approximately 2.24 times slower than cells cultured at 37°C (Fig. 2i). These results demonstrated that the DivisionCounter effectively detects and quantifies variations in division rates, showcasing its utility for quantifying modulation of division rate across conditions by simply measuring FP expression loss.

In conclusion, we validated and benchmarked the DivisionCounter method, demonstrating its utility and accuracy to measure division counts and division rates. The DivisionCounter allows up to 74 divisions to be robustly counted, surpassing the 10-division limit of other state-of-art methods and the Hayflick limit of 40-60 divisions.

### DivisionCounter measures B-ALL cell division rates across organs *in vivo*

To assess the DivisionCounter’s performance *in vivo*, we employed the well-established Nalm6 leukemia xenograft model in immunodeficient NSG mice. In this model of tumour development, Nalm6 cells progressively grow and spread across various organs, with initial growth in the bone marrow (BM) and liver, then in the spleen and lungs^22,23^. While different organs display different kinetics of tumour invasion, it is unclear whether these distinct environments directly impact Nalm6 growth. To investigate the effects of the organ microenvironments on Nalm6 cell divisions, we applied the DivisionCounter method within this *in vivo* tumour model.

We first validated *in vitro* the use of the 26G-DivisionCounter on mTagBFP2^+^luciferase^+^ Nalm6 cells. After infection, Nalm6 cells were cultured, counted and assessed for the percentage of mScarlet-positive cells, for up to 59 days. As expected, the percentage of mScarlet-positive Nalm6 decreased with the cell population doubling and reached the 1/3 equilibrium (Extended Data Fig. S3a). In contrast, in the 26N-control–infected Nalm6 cell culture, the percentage of mScarlet-positive cells remained stable over time (Extended Data Fig. S3a). This validated the use of the DivisionCounter in Nalm6 cells. Fitting a nonlinear-mixed-effects model to the loss of percentage of mScarlet-positive cells over the average cell population doubling, we obtained a probability of losing mScarlet-expression in Nalm6 cells of 0.049 (CI= 0.046-0.057, Extended Data Fig. S3a), similar to the HEK293T cells value (Fig. 2c).

After injecting 2.5 x 10^5^ Nalm6 cells infected with the 26G-DivisionCounter into NSG mice, we followed Nalm6 tumour development over time (at days 7, 13 and 20 after *in vivo* injection) using the DivisionCounter and standard measurements of *in vivo* cell growth (luciferase assay, KI67 staining, and cell counting) (Fig. 3a). Luciferase measurements showed a clear exponential growth of Nalm6 throughout the whole body but reached saturation at day 20 (Fig. 3b and Extended Data Fig. S3b), preventing any further measurement of Nalm6 total body growth. The Ki67 measurements indicated that most cells were dividing (Ki67+) in all organs, with no significant differences between them (Extended Data Fig. S3c). None of these methods were informative with respect to whether the distinct organ microenvironments affect Nalm6 growth, illustrating their limitations. Nalm6 cell counts measured by flow cytometry with counting beads showed a higher number of cells in the BM and liver than in the lungs, blood or spleen across all time points (Fig. 3c, Extended Table S2). These results were consistent with previously published kinetics of tumour invasion^22,23^. These differences in cell counts indicated higher net cell growth in the BM and liver as compared to the other organs. However, it was impossible to conclude if these differences in Nalm6 cell counts were due to faster division rate, delayed or lower initial cell seeding, or differences in migration between organs.

**Figure 3:**
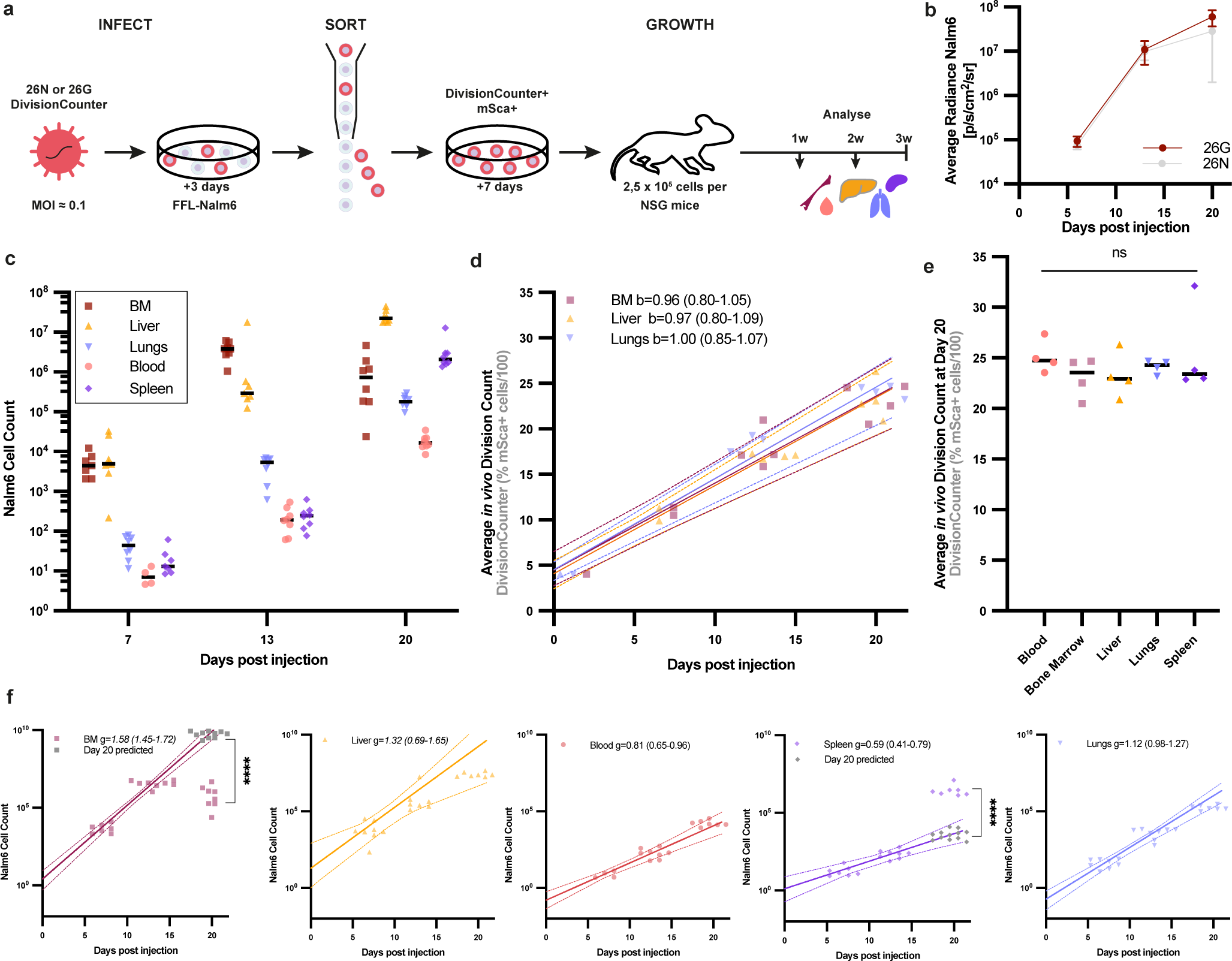
the DivisionCounter measures the *in vivo* average division count and rate of Nalm6 leukemia cells. a) mTagBFP2^+^Luciferase^+^Nalm6 cells were infected with the 26G- or 26N-DivisionCounter lentivirus at a low multiplicity of infection. After 3 days of culture, mScarlet-positive cells were sorted injected and cultured for 7 days before 2.5 x10^5^ 26G-DivisionCounter infected Nalm6 cells were intravenously injected in NSG mice. Nalm6 tumour development was then followed over time by isolating cells at day 7, 13 and 20 after *in vivo* injection from the blood, bone marrow (BM), spleen, liver and lungs. b) average radiance of Luciferase measurement in the 26G-DivisionCounter (red) and the 26N-DivisionCounter (grey) infected Nalm6 cells at day 7, 13 and 20 after *in vivo* injection. c) Nalm6 cell counts (CD45^-^HLA^+^) over time in blood, spleen, lungs, liver and BM. Each dot is one mouse, n=7, 8, 7, 4, 6 mice at day 7 for BM, liver, lungs, blood, spleen respectively and n= 8 mice at day 13 except for Liver (n=7), the 26G and 26N- condition were pooled. d) DivisionCounter average division counts computed using the probability from S3a over time in the BM, liver and lungs. Each dot is mouse, n=4 mice per time points, except for BM at day 7 n=2, liver at day 7 n=2, lungs at day 13 n=3, measurements were done in triplicates. A linear regression was fitted to the average division count over time, b= the obtained division rate with CI computed using bootstrapping. The lines represent the average and CI for the expected division count using the fitted division rates. e) DivisionCounter average division count computed using the probability from S3a at day 20 in blood, spleen, lungs, liver and BM. Average value over triplicate per mice, n=4 mice. Statistical comparison done with permutation testing. f) Nalm6 cell counts (CD45^-^HLA^+^) over time in blood, spleen, lungs, liver and BM as in c. An exponential regression was fitted to the cell counts with the obtained average net cell growth rate (g) value indicated for each organ with CI computed using bootstrapping. The lines represent the average and CI for the expected cell count using the fitted net growth rates. The grey dots are the points predicted using the fitted exponential growth for the BM and the Spleen. Each dot is a mouse, n=8 mice per time points with the 26G and 26N- condition pooled.

We next tested the DivisionCounter method to measure Nalm6 proliferation *in vivo*. To first assess the usability and robustness of the DivisionCounter method in measuring the average division count of Nalm6 cells *in vivo*, we used the 26G-DivisionCounter and assessed the percentage of mScarlet-positive cells at days 7, 13 and 20 after injection (Fig. 3d). We validated the use of the DivisionCounter *in vivo* by demonstrating a robust mScarlet signal after *in vivo* injection (Extended Data Fig. S3d) and a stable percentage of mScarlet-positive cells over time in the 26N-control Nalm6 cells (Extended Data Fig. S3e). In addition, the DivisionCounter did not impact the growth properties of Nalm6 cells, as no statistical differences in cell growth between 26G-DivisionCounter and 26N-control infected Nalm6 cells were observed, based on both luciferase and cell count measurements (Extended Table S3). Next, we calculated the DivisionCounter average division count of Nalm6 by inputting the percentage of mScarlet-positive cells with the 26G-DivisionCounter into the DivisionCounter formula. This revealed that, in all organs and at all time points, Nalm6 cells had divided more than 10 times (Fig. 3d). The DivisionCounter results were robust and highly sensitive, with limited variability between mice even when as few as 455 Nalm6 cells were measured (Extended Data Fig. S3f). Using the DivisionCounter, we determined that Nalm6 cells had divided ∼11, 18, 23 times at days 7, 13 and 20 post injection, respectively, allowing for the first time to measure the division counts of tumour *in* vivo. These values could not have been measured by any existing state-of-the-art method. Overall, the DivisionCounter method demonstrated high sensitivity and robustness for *in vivo* counting of divisions.

To evaluate the impact of the organ microenvironment on Nalm6 cell division, we compared the *in vivo* average division counts obtained with the DivisionCounter in the different organs. The average division counts were similar between the bone marrow (BM), liver and lungs at days 7 and 13 (Fig. 3d). Similarly, by day 20, when the Nalm6 had spread well into all organs, the DivisionCounter measured on average 21-26 divisions in all organs (Fig. 3e), with no statistical differences between blood, spleen, BM, liver or lungs (Extended Table S4). These results indicated that the microenvironment provided by different organs did not impact Nalm6 division. This conclusion was further confirmed by fitting a linear regression to the division count over time: the DivisionCounter method revealed that Nalm6 cells in BM, liver and lungs all divided at a rate of ∼1 division/day (Fig 3d and Extended Table S5), with no statistical difference between these three organs (Extended Table S6). Thus, the DivisionCounter demonstrated that the *in vivo* division rates of Nalm6 leukemia tumor cells in NSG mice were not modulated by the organ-specific microenvironment.

We next integrated the DivisionCounter results with cell counts to gain further biological insight into the kinetics of tumour invasion in different organs. Between the days 13 and 20, Nalm6 cell counts stopped increasing in the BM but continued to increase in the other organs (Fig. 3c). Indeed, the cell counts in the BM were significantly lower than expected by exponential growth (Fig. 3f and Extended Table S7-8). The plateauing of Nalm6 cell counts in the BM could be due to a cessation of cell division or to increased cell death, but cell counts alone cannot resolve those hypotheses. The DivisionCounter, however, revealed that Nalm6 cells in the BM continued to divide at the same rate between days 13 and 20, with no change in the percentage of dead cells observed at day 13 and 20 (Extended Data Fig. S3g), demonstrating that the lower number of Nalm6 cells in the BM was not due to change in division or death. An alternative hypothesis is that the tumour cells lacked space in the BM and egressed to compensate for their growth, as observed in an acute myeloid leukemia mouse model ^24^. In support of this hypothesis, we observed a significant increase in Nalm6 cell counts in the spleen at day 20, higher than expected due to exponential growth (Fig. 3f and Extended Table S7-8), along with the presence of Nalm6 in the blood (Fig. 3c), both suggesting increased circulation of Nalm6 cells between organs. Therefore, the stable Nalm6 cell counts in the BM would be well explained by a BM space saturation coupled with the migration of Nalm6 cells into the circulation and spleen. Additionally, we observed a lower Nalm6 cell counts in the lungs at days 7 and 13 as compared to BM and liver (Fig. 3c). As the DivisionCounter method showed that Nalm6 cells had the same average division counts at days 7, 13 and 20 in lungs, BM and liver (Fig. 3d), with no significant difference in division rates between the lungs and liver at day 20 (Fig. 3e), the differences in division rate were not responsible for the lower cell count in the lungs. It is more likely that the lower Nalm6 cell counts in the lung were due to a lower number of Nalm6 cells seeding the lungs, or to a later tumour seeding in the lungs from Nalm6 cells migrating from the BM or liver via the circulation. Therefore, the DivisionCounter is more informative than cell counting alone, and using the DivisionCounter in conjunction with cell counting significantly enhanced our understanding of the kinetics and spatial dynamics of Nalm6 tumor development *in vivo* across multiple organs.

### DivisionCounter measures the *in vivo* tumour division rate during CAR-T cell therapy, allowing the CAR-T cell *in vivo* killing rate to be inferred

Next, we investigated the utility of the DivisionCounter in deconvolving the effect of cell division and cell death in the context of targeted tumour killing cell therapies. We explored *in vivo* how a targeted killing cell therapy impacted the Nalm6 B-ALL cell proliferation using the DivisionCounter method. Adoptive cell therapies with T cells expressing chimeric antigenic receptor (CAR) have efficient clinical results against B-ALL cells^25^. CAR-T cells efficiently kill tumour cells either directly or via the microenvironment^26,27^ but it is unknown if they directly impact tumour cell division. To evaluate this, we used the DivisionCounter to quantify the average division counts over time in the presence or absence of CAR-T cell therapy.

Seven days after injecting 2.5 x 10^5^ 26G-DivisionCounter–infected Nalm6 cells into NSG mice, we transferred 2 x 10^6^ T cells transduced with the 4-1BB-based CD19 CAR construct ^28^ or mock construct into separate mice (Fig. 4a). Luciferase measurements revealed a lower average Nalm6 radiance, suggesting a lower Nalm6 tumour burden at day 13 days post-injection (DPI) with T-cells, in CAR-T cell-treated mice as compared to the mock-T cell-treated mice (Fig. 4b). This reduced tumour burden could already be observed in the BM at 7 DPI (Fig. 4c), whereby BM Nalm6 cell counts and Nalm6 net cell growth in the CAR-T cell-treated group were significantly lower than in the mock-T cells group (Extended Table S9). This decrease in Nalm6 net cell growth was due to CAR-T cell killing of Nalm6 cells. Indeed, CAR-T cells expanded *in vivo* in response to Nalm6, with CD3+ T cell counts significantly higher in the BM of CAR-T cell treated mice than mock-T cell-treated mice at 7 DPI with T cells (Extended Data Fig. S4a-b). Additionally, CAR-T treatment induced more Nalm6 cell lysis than mock-T cell treatment, as measured by an *in vitro* cytotoxic assay, with no differences observed in killing of the Nalm6 cells with 26G-DivisionCounter or 26N-control infection (Extended Data Fig. S4c-d). Fitting an exponential growth model onto cell count data, we found a net growth rate of the Nalm6 cells of 0.77 cells/day in BM of the CAR-T cell treated group, which is lower than the 1 cell/day growth rate of the mock-T cell group (Fig. 4c). Thus, CAR-T cells were clearly killing Nalm6 and inducing a reduced Nalm6 cell counts over time, yet it was still unclear whether CAR-T cells affected Nalm6 divisions.

**Figure 4:**
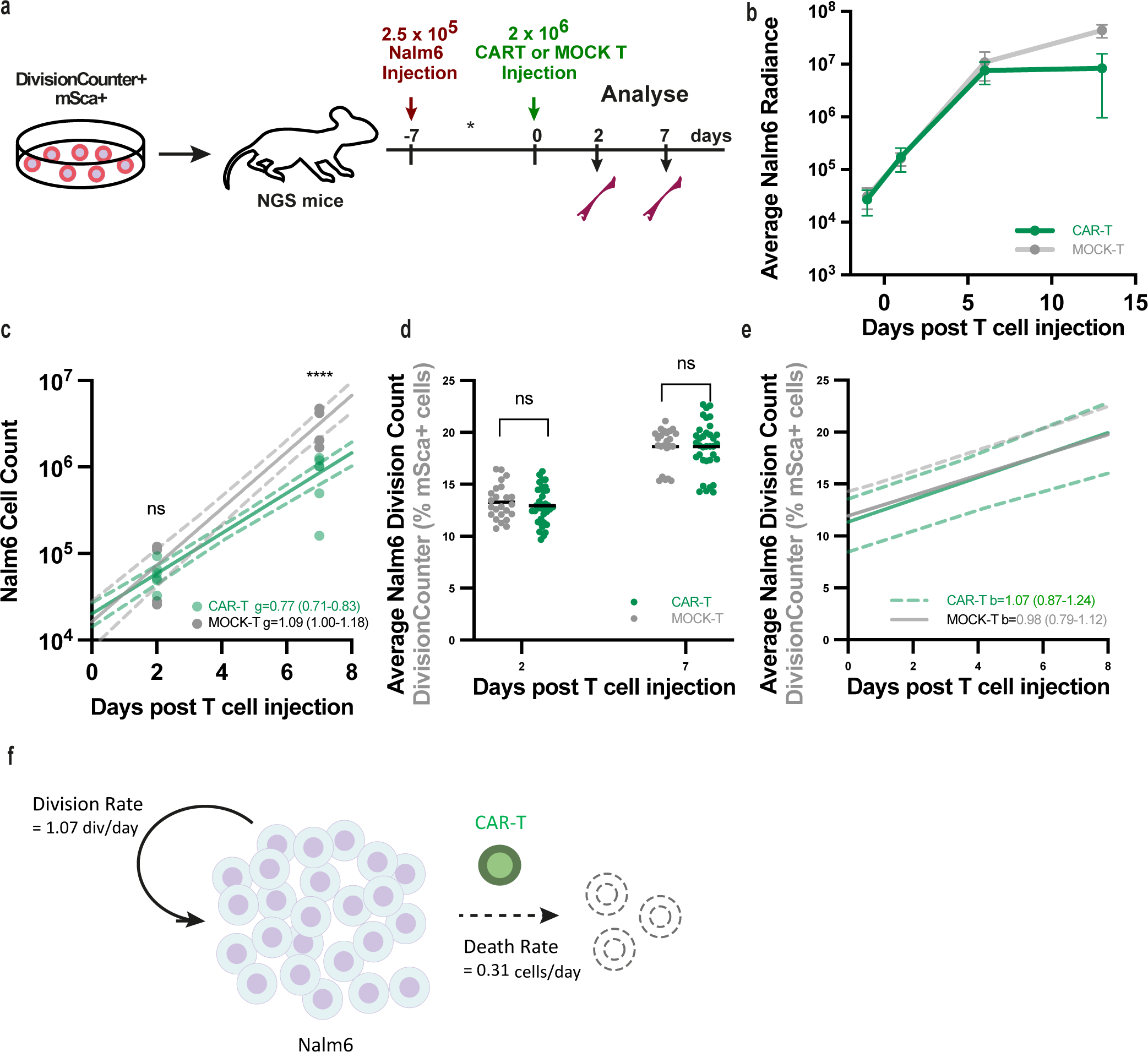
the DivisionCounter measured the division count and rate of Nalm6 leukemia cells upon CAR-T treatment. a) mTagBFP2^+^Luciferase^+^Nalm6 cells were infected with the 26G-DivisionCounter lentivirus at a low multiplicity of infection. After 3 days of culture, mScarlet-positive cells were sorted injected and cultured for 7 days. 2.5 x10^5^ 26G-DivisionCounter infected Nalm6 cells were injected into NSG mice. 7 days later 2 x10^6^ 4-1BB CAR-T or Mock-T were injected. Tumour development was followed over time by isolating cells from the BM at day 2 and 7, 13 post T cell injection. b) average radiance of Luciferase measurement mice injected with CAR-T (green) or Mock-T (grey) at day 2 and 7 post T cell injection, n= 12 mice, error bars display the 95% CI. c) Absolute Nalm6 cell counts (CD45^-^HLA^+^) over time in the BM of mice injected with CAR-T (green) or Mock-T (grey). An exponential regression was fitted to the cell counts with the obtained average net growth rate (g) value indicated for each condition with CI computed using bootstrapping. The lines represent the average and CI for the expected cell count using the fitted net growth rates. Each dot is one replicates, 10 replicates per mouse, n=6 mice per time points for the CAR-T condition and 4 per time points for the Mock-T condition. ns indicates non significance and *** indicates p<0.001 using permutation testing. d) Average division count of 26G-DivisionCounter Nalm6 cells computed using the DivisionCounter formula based on the percentage of mScarlet-positive cells analysed by flow cytometry at each time point in the BM of mice injected with CAR-T (green) or Mock-T (grey). Each dot is an experimental data point. N=6 mice per time points for the CAR-T condition and 4 per time points for the Mock-T condition. ns indicates non significance using permutation testing. e) Fitted division rate of 26G-DivisionCounter Nalm6 cells over time based on the average division count in d) in the BM of mice injected with CAR-T (green) or Mock-T (grey). b is fitted division rate value with CI computed using bootstrapping. The lines represent the average and CI for the expected division counts using the fitted division rates. f) Schematic of the Nalm6 division rate and CAR-T killing rate.

We next used our DivisionCounter method to assess whether the reduced Nalm6 net cell growth in the CAR-T cell treated group could be attributed solely to CAR-T cell killing of Nalm6 or if changes in Nalm6 division rate occurred *in vivo*. First, by inputting the percentage of mScarlet-positive cells observed in the BM over time into the DivisionCounter formula, we quantified that Nalm6 tumour cells had divided over 10 times *in vivo* at 2 DPI and 7 DPI of T cells (Fig. 4d and Extended Data Fig. S4a), demonstrating the necessity of the DivisionCounter method for counting over 10 divisions in therapeutic contexts. Importantly, this quantification revealed no statistical differences in Nalm6 division counts between the CAR-T cell and mock-T cell treated mice (Fig. 4d), demonstrating that Nalm6 division was not affected by CAR-T cell killing. Additionally, Nalm6 cells divided at 1 division/day (Fig. 4e), comparable to the rate without CAR-T injection (Fig. 3d), further supporting the finding that the Nalm6 division rate was not impacted by the presence or actions of CAR-T cells. Lastly, combining the DivisionCounter inferred division rate with the cell count inferred net cell growth rate, we estimated the Nalm6 cell death rate induced by CAR-T cells to be on average 0.31 cell/day (CI= 0.21-0.43, Fig. 4f). This low value could be due to the sub-optimal number or potency of the CAR-T cells, demonstrating the power of the method to evaluate the factors influencing tumour growth during treatment.

In conclusion, the DivisionCounter is a robust method for quantifying cell division *in vivo* in animal models, in a cell population subject to complex cell population dynamics that include death and migration. The DivisionCounter is the first method capable to quantify cell division and death rates *in vivo* over large division ranges, enabling the user to quantify the contributions of division and other factors—such as death and migration—to cell growth. We have showcased the DivisionCounter’s utility to decipher the kinetics of tumor growth and effect of tumour therapies.

## Discussion

In this study, we developed the DivisionCounter method, a robust, accurate and versatile cell population division counter that can quantify cell divisions and division rate over time and in different conditions. The DivisionCounter method comprises a genetic construct and a mathematical formula, and it outperformed start-of-the-art methods by counting between 56 and 74 divisions for its 26G- and 23G-versions, respectively, reaching the Hayflick limit of 40-60 divisions^1^ and the estimated division number of most cells. Indeed, 44 divisions without death would produce all 10^13^ cells in a human body^29,30^.

The chief assets of the DivisionCounter include: *i*) a quantitative framework comprising the DivisionCounter formula and statistical testing to estimate confidence interval and division rate over time; *ii*) a well-defined working range and accuracy. The DivisionCounter counts divisions until it reaches the equilibrium between the three states of its genetic construct and, after this point, still provides a lower bound of division count, i.e. that cells have undergone at least this division count to reach the 1/3 equilibrium; *iii*) a tuneable working range of division counting by changing the motif or length of the microsatellite in the DivisionCounter construct; and *iv*) an easy readout using fluorescence by flow cytometry and microscopy with robust values for as little as 455 cells analysed.

We have demonstrated the utility of the DivisionCounter method for quantifying division counts and division rate *in vivo*, consequently allowing death rates to be estimated *in vivo*. Changes in net cell growth rate due to microenvironment, treatments or ageing have been traditionally assessed using cell counting, division tracker dyes (e.g. CTV, CFSE), snapshot measurement of cell cycle (e.g. Ki67) and death markers (e.g. Caspase). In contrast to these methods, the DivisionCounter allows the *in vivo* division rate to be deconvoluted from the death and migration rates. This offers the potential to quantify systemic effects on growth rate with applications in CAR-T cell development, as well as drug and genetic screenings to evaluate drugs or gene knock-down effects on cell division or death.

The DivisionCounter method offers a broad applicability. For instance, it could be reengineered and expanded to non-mammalian cells, or it could be used with single-cell omic readouts to compute the average division count within cluster of cells. Additionally, more than one DivisionCounter construct could be used, each with a distinct microsatellite and FP, to provide multiple ranges of division count resolution. For cells that are difficult to transduce using lentiviruses, it would be possible to adapt the DivisionCounter method to include a genetic mouse model, similar to the ones used in mosaicism research^31^. Overall, the DivisionCounter method provides the potential to study the role of cell division in healthy and pathological tissue development *in vitro* and *in vivo*.

## Material and Method

### DivisionCounter construct cloning

To construct the DivisionCounter backbone, the pLVX-EF1α-AcGFP1-N1 plasmid (Clontech 631983) was amplified using primers targeting *EcoR1* restriction site and the start of *puroR* gene, in order to remove *AcGFP1* and *PGK* promotor from the plasmid. *mScarlet* gene without start codon followed by a P2A element upstream of puroR was added using pmScarlet_C1 (Addgene #85042) plasmid as template. A two-site directed mutagenesis to insert a *Nde1* restriction site downstream puroR and a *Spe1* restriction site upstream of *EcoR1* was performed. In the end we obtained the EF1alpa-Spe1-EcoR1-mScarlet-P2A-puroR-Nde1-WPRE plasmid, that was used as our reference DivisionCounter backbone.

Small (≤ 100bp) oligonucleotides were synthetized by Eurofins, of the form Spe1-Kozak-Microsatellite-EcoR1, where the Microsatellite was either a guanine repeat (23G, 26G) or a spacer sequence made of 26 nucleotides (26N). All oligonucleotide sequences are supplied in table X. Those small oligonucleotides were hybridized and subsequently ligated into the DivisionCounter backbone previously digested by Spe1(NEB) and EcoR1(NEB) using T4 ligase (NEB). The resulting ligation reaction was added to JM109 competent bacteria that were subsequently grown in SOC medium before being plated onto agar plate containing ampicillin and incubated overnight at 30°C. Clones of interest were then selected, amplified, sanger sequenced to verify the correct ligation of our oligomers and one of those clones was then subsequently amplified in maxi preps to generate our final plasmid stock. All oligos used are in Table S10.

### Lentivirus production, transduction and titration

HEK293T cells were plated at seeding density of 12x10^6^ in T150 in DMEM glutamax (Gibco 61965-026) supplemented with 10% FCS (Dutscher S1900-500C), 1% sodium pyruvate (Gibco, 11360070) and 1% MEM-NEAA(Gibco, 11140050) and grown overnight at 37°C with 5% CO2. The following day, the medium was replaced, and cells were transfected with the DivisionCounter plasmid, the psPAX2 lentiviral packaging plasmid (Addgene plasmid #12260,) and the pMD2.G envelope plasmid (Addgene, plasmid #12259). Using a 2.5:1 ratio of psPAX2:pMD2G, the packaging and envelope plasmids were incubated in a NaCl solution (150 mM) containing 1:12 Polyethylenimine “Max” (PEI MAX; polyScience #24765, 7.5 mM) for 15 minutes. Two days after transfection, supernatant was collected and filtered through a 0.45 µm filter and concentrated using Amicon Ultra-15 filters (100 kDa) (Millipore #UFC9100) via centrifugation. Viruses were then eluted in Opti-MEM (Gibco, 31985062) and stored at -80 °C until further use.

The viral titter and concentration were determined for each cell line by titrating the viral stock on 5x10^5^ newly plated cells (at 0.5 × 10^5^ cells/mL in a 24-wells plate) in their specific culture medium and measuring mScarlet expression 3 days post infection by flow cytometry with Cytoflex LX (Beckman). A MOI of 0.1 was targeted to ensure the majority (over 90%) of the infected cells would have been successfully transduced with only one construct. DivisionCounter experiment *in vitro*

### Culture medium and DivisionCounter infection of HEK293T cells

Human embryonic kidney 293 T (HEK293T) cells were cultured at 37°C, 5% CO2 in Dulbecco’s Modified Eagle’s Medium (DMEM+GlutaMAX; Gibco 61965-026), supplemented with 10% heat inactivated Fetal Calf Serum (FCS; Dutscher S1900-500C) and 1% Penicillin Streptomycin (Pen Strep; Gibco 15140-122).

HEK293T were transduced with DivisionCounter viruses at MOI 0.1. 2 to 6 days later, HEK293T cells were collected, stained with SYTOX Green (Invitrogen, R37168, 1/40 dilution, incubated no more than 30 min before acquisition) and live mScarlet-positive cells were sorted on BD ARIA FUSION (BD Bioscience). 25000 sorted mScarlet+ HEK293T were subsequently plated in a 24-wells plate and incubated at 37°C or 33°C with 5% CO2. Cells were split twice a week for up to 46 days post sort by removing the cells supernatant, incubating with TrypLE Express (Gibco, 12605010) at 37°C, 5% CO2 before resuspending in full medium, counted and plated back in a fresh plate or flask.

### Average Cell Division Count from cell count measurement (cell population doubling)

At each acquisition time point, before each splitting, we counted the absolute number of cells in each well using the Scepter 3.0 handheld automated Cell counter (Merck, PHCC360KIT, PHCC360250). To obtain the cell population division count over time, we computed the cell population doublings (PD) at each time point (*t*) using the absolute cell counts :

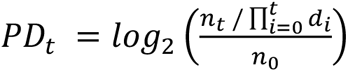

where nt is the absolute number of cells measured at each time point, di is the inverse of the cumulative product of splitting ratios, and n0 is the number of cells initially plated. Assuming cells grew exponentially with no cell death; the cell population doubling then equals the average division count of the cell population.

### m-scarlet measurements for the DivisionCounter read-out on HEK293T

When splitting HEK293T cells, left over HEK293T cells were washed and stained with SYTOX Green Green (Invitrogen, R37168, 1/40 dilution, incubated no more than 30 min before acquisition). mScarlet expression was measured with Cytoflex LX (Beckman) and the percentage of mScarlet-positive cells over time were analysed with FlowJo (BD Bioscience) as in Figure S2a. This percentage was used either for the statistical inference of the fluorescence loss probability or to compute the average division count with the DivisionCounter formula (see below).

### mScarlet reacquisition experiment with HEK293T

In order to track mScarlet reacquisition in HEK293T, we followed a similar protocol as detailed before except that we sorted mScarlet-negative cells from sorted mScarlet-positive cells that were left in culture for 20 to 40 days post sort and that had lost their mScarlet expression in the meantime/as they divided. Cytometry analysis, cell counting and cell population doubling computation were performed as before.

### In vitro experiments on other cell types (MDA-MB-468, MEF, and Nalm6)

We performed similar *in vitro* experiments as with HEK293T cells on Mouse Embryonic Fibroblast (MEF), Human MD Anderson-Metastatic Breast-468 (MDA-MB-468), and Nalm6 cells. MEF were cultured in the same medium as HEK293T cells, with inactivated FCS (Gibco 15140-122). MDA-MB-468 cells were cultured in Dulbecco’s Modified Eagle’s Medium (DMEM+GlutaMAX; Gibco 61965-026), supplemented with 10% heat inactivated One Shot Fetal Bovine Serum (Gibco A3840401). Human Nalm6 cells, a B cell precursor leukaemia cell line, expressing firefly luciferase (Fluc) and blue fluorescent protein (BFP), called Nalm6 in this paper, were cultured in Roswell Park Memorial Institute (RPMI) 1640 Medium with GlutaMAX™ Supplement (Gibco, 61870-010), supplemented with 10% heat inactivated Fetal Calf Serum (FCS; Dutscher S1900-500C, Batch: S00CH) and 1% Penicillin Streptomycin (Pen Strep; Gibco 15140-122) at 37°C, 5% CO2.

After transduction and sorting of mScarlet+ cells, MDA-MB-468 and MEF were plated at 5x10^4^ cells/ml seeding density and Nalm6 cells at 10^6^ cells/ml seeding density and cultured all at 37°C, 5% CO2. Every 2-4 day, cells were split and, at each acquisition time point, cells were replated at 10^5^ cells/mL for MDA-MB-468 and MEF and between 1-3x10^6^ cells/mL for Nalm6. Nalm6 cells being more sensitive to sort, we sorted them at room temperature and collected them in pure Fetal Calf Serum (FCS; Dutscher S1900-500C, Batch: S00CH) before replating them in culture medium.

When splitting MEF, MDA-MB-468 and Nalm6 cells, left over cells were washed and stained with SYTOX Green Green (Invitrogen, R37168, 1/40 dilution, incubated no more than 30 min before acquisition). mScarlet expression was measured with Cytoflex LX (Beckman) and the percentage of m-Scarlet-positive cells over time were analysed with FlowJo (BD Bioscience) as for HEK293T cells.

### DivisionCounter experiment in vivo Mice

All mouse studies were carried out in accordance with guidelines and approval of French authorities. All *in vivo* experiments were conducted using 8- to 16-week-old NOD/SCID/IL-2Rγ-null (NSG) female mice purchased from Charles River Laboratories (France), housed in a facility with 12-h light/12-h dark cycle at 22 ± 1 °C and 40 ± 10% humidity. Teklad Global 18% protein rodent diet and tap water were provided ad libitum. All mouse experiments were performed using protocols approved by the French Ministry of Higher Education, Research and Innovation. Ethics committee number 118 (APAFIS#26128-2019032323036339, DAP2018-014 and APAFIS#28212-2020111714259701, DAP2020-021)

### CAR-T cell production

Buffy coats from anonymous healthy donors were obtained from Etablissement Français du Sang (Paris, France) in accordance with INSERM ethical guidelines. According to French Public Health Law (art L 1121-1-1 and art L 1121-1-2), written consent and Institutional Review Board (IRB) approval are not required for human noninterventional studies. Peripheral blood mononuclear cells (PBMCs) were isolated from healthy donor leukapheresis rings by density gradient centrifugation (Lymphoprep, Stemcell 07851) and frozen in CryoStor cell cryopreservation media (Sigma-Aldrich C2874). T lymphocytes were purified using the Pan T cell isolation kit (Miltenyi Biotech 130-096-535) and activated with Dynabeads human T-Activator CD3/CD28 (1:1 bead:cell) (ThermoFisher 11131D) in X-VIVO 15 medium (Lonza 02-053Q) supplemented with 5% human serum (Sigma Aldrich H4522) and 0.5 mM b-mercaptoethanol at density of 10^6^ cells/mL. 24 hours after activation, T cells were transduced with lentiviral supernatants of an anti-CD19 (FMC63)-CD8tm-4IBB-CD3ζ CAR construct (rLV.EF1.19BBz, Flash Therapeutics) at MOI 10 in complete X-VIVO with a final concentration of 4 μg/mL of Polybrene. After 16-18 hours the supernatant was almost completely discarded, and cell suspension was completed to a final volume of 2 mL with complete X-VIVO containing 100 IU/ml of rhIL-2 (Peprotech 200-02). For untransduced T cells, the same protocol was followed without lentiviral particles. CAR-T cells were maintained in X-VIVO supplemented with 100 IU/ml of rhIL-2 and 6 days after transduction, cells were subjected to magnetic removal of activation beads and collected to check CAR expression, phenotyping, killing capacity or freezed for later injections into tumor-bearing mice.

CAR-T cells phenotyping was assessed using a CD19 CAR detection reagent (Biotinylated) (Miltenyi Biotech 130-129-550) and incubating for 10 minutes at room temperature. After washing, cells were stained with PE-conjugated streptavidin for 30 minutes at 4°C in PBS with BSA 0.5% and EDTA 2 mM. Samples were acquired on BD LSR-II or BD Fortessa X-20 (BD Biosciences) and analyzed with FlowJo software v.10.6.2 (BD Biosciences).

### Cytotoxicity assays

The cytotoxicity of T cells transduced with a CAR was determined by co-culturing in triplicates at the indicated *E*/*T* ratio, CAR-T cells (effector cells) with NALM6 Luc and BFP-expressing, 26G-DivisionCounter or 26N-DivisionCounter, (target cells) in a total volume of 100 μl of complete X-VIVO per well using U bottom 96-well plate. Target cells were plated at the same cell density alone to determine the maximal luciferase expression (relative light units (RLU)). Effector cells were plated alone to measure background. 18 h after culturing the cells at 37°C, 50 μl of luciferase substrate (Perkin Elmer 122799-10) at 0.3 mg/ml in PBS were added to each well and luminescence was detected in a SpectraMax ID3 plate reader (Molecular Devices). % Specific Lysis was determined as (1 - (RLUsample - RLUbackground)/(RLUmax-RLUbackground)) x 100.

### Nalm6 infection and sorting

0.25x10^6^ NALM6 BFP-expressing 26G-DivisionCounter cells were injected into 8–16-week-old female NSG mice intravenously by tail vein injection. Three days later 2x10^6^ CAR-T cells were thawed and injected via lateral tail vein, defining the day 0 of the experiment. Tumour burden was measured by bioluminescence imaging using Xenogen IVIS Imaging System (PerkinElmer) one or two times per week. Acquired bioluminescence data was analysed using Living Image software (PerkinElmer) and expressed in average radiance (photons/sec/cm2706 /sr).

### In vivo cell injection

0.25x10^6^ Nalm6 cells were injected into NSG female mice intravenously via lateral tail vein injection. For the CAR-T cells experiment, seven days later 2x10^6^ CAR-T cells were unfreezed and injected via lateral tail vein, defining the day 0 of the experiment.

### In vivo Luciferase assay

Nalm6 tumor burden was measured by bioluminescence imaging using Xenogen IVIS Imaging System (PerkinElmer) one or two times per week. Acquired bioluminescence data was analysed using Living Image software (PerkinElmer) and expressed in average radiance (photons/sec/cm2706 /sr).

### Mouse tissue processing and cell isolation

At each acquisition time point, blood was first collected from the submandibular vein from each mouse. Then cervical dislocation was performed and both femurs and tibias, lungs, liver, and spleen were collected for each mouse and all organs were processed to obtain cell suspensions before antibody staining.

For each blood sample (200uL), red blood cells (RBC) were lysed by hypotonic shock with distilled water and filtered using 70µm cell strainer before flow cytometry staining.

Bone marrow cells were isolated from femur and tibia bones of mice by bone flushing for (Fig 3) or centrifugation (300gx 30sec-1min) for (Fig 4). All BM cells were then washed and stained for flow cytometry analysis, except BM cells from the day 7 of the figure 3 experiment that were first enriched. For the enrichment, cells were filtrated through a 70µm cell stainer and washed in cold MACS buffer (PBS-0,5% BSA). Up to 140.10^6^ cells were resuspended in 80µl of MACS buffer and incubated 15min at 4°C with 20µl of Mouse cell depletion Cocktail (Miltenyi 130-104-694). Prior magnetic separation cells concentration was adjusted to 30.10^6^ cells/ml using MACS buffer. Enrichment was done with LS column (Miltenyi #130-042-401) following the manufacturer recommendations. Cells were then washed twice with PBS supplemented with 10%FBS then stained for flow cytometry analysis as the other BM cells.

Spleen cells were isolated by crushing spleen through 100µm nylon mesh cell strainer (fisherbrand 22363549) with the plunger end of a syringe, then washed and stained for flow cytometry analysis according to the antibody panel presented below.

Lungs were cut in small pieces and digested with Liberase TL ((Roche, ref: 05401020001, 0.15 mg/mL) and DNAse (Roche, ref: 11284932001, 1 mg/mL) at 37°C for 30 minutes. With the plunger end of a syringe, digested lungs were mashed through a 100 μm cell strainer and washed to obtain single cell suspensions. Before staining, mononuclear cells were enriched using a density gradient medium (Lymphoprep, STEMCELL technologies, Catalog#07851) and centrifugation at 400xg for 30min at room temperature.

Livers were injected and incubated for 15 minutes at 37°C with dissociation buffer, dPBS supplemented with 0,01 mg/ml Collagenase D (Roche 11088858001) and 200U/ml DNAse I (Roche 10104159001), then dissociated using GentleMACS (Miltenyi 130-096-427) 37C_m_LIDK_1 programme. Before staining, mononuclear cells were enriched using a density gradient medium (Lymphoprep, STEMCELL technologies, Catalog#07851) and centrifugation at 400xg for 30min at room temperature After cell isolation, cells were either directly stained and acquired (figure 3) or kept in CO2-independant media (Gibco 18045-054) overnight before staining and acquisition (figure 4). All cells were stained as explained below.

### Antibody staining and flow cytometry acquisition and analysis

After cell counting using the LUNA-FX7™ (Logos biosystems), the total number of cells per sample was distributed into 1 to 10 wells with a maximum of 2x10^6^ cells/well. In at least 3 wells/sample 50ul CountBright^TM^ absolute counting beads (ThermoFisher C36950) were added. Cells were then stained with mCD45, mTer119, hCD19, hHLA-DR for Fig 3), and also hCD3-BV510 (for Fig 4) antibodies (details in table S11) in PBS with 10% FBS at 4°C for 20 min in the dark. The live/dead cell marker (SYTOX™ Green Ready Flow™ Reagent Invitrogen™ R37168) at a 1/40 dilution was added no more than 30min at RT before acquisition.

For BM enriched samples and Ki67 staining samples (Fig 3), cells were washed twice in PBS then stained with Live/Dead (LIVE/DEAD™ Fixable Near-IR Dead Cell Stain Kit Invitrogen™ L3224) at 1:100 dilution for a maximum of ∼10^6^ cells/100ul for 15min at room temperature in the dark. Cells were then washed in PBS with 10%FBS and incubated with hCD19 and hHLA A,B,C antibodies (details in table S11) in PBS with 10% FBS at 4°C for 20 min in the dark. The cells were then washed in PBS with 10%FBS and either brought for acquisition for the BM enriched samples, or fixed and stained for Ki67 staining samples.

Ki67 staining samples were fixed and permeabilized using the FOXP3 Fix/Perm Buffer Set (Biolegend 421403) following the manufacturer recommendations. Ki-67 antibody staining was then performed in Permeabilization Buffer (from the set) at 4°C for 30minutes, before cells were washed and resuspended in PBS with 10%FBS.

Flow cytometry data were acquired on Cytoflex LX (Beckman) and analysed in FlowJo (BD Bioscience). Nalm6 cells were gated as hHLA-DR^+^ cells in figure S3d and S4a except for day 20 data of Figure 3, for which a hHLA-DR^+^hCD19^+^ was used. mScarlet-expression was gated as in figure S3d and S4a and the percentage of mScarlet-positive cells percentage was used for analysis using the DivisionCounter mathematical framework (see below). T-cells were gated as in Figure S4a.

### Absolute Cell count per sample using CountBright^TM^ absolute counting beads

To retrieve absolute Nalm6 and T cell counts per sample, we used CountBright^TM^ absolute counting beads as per manufacturer recommendation. A fixed volume (50µl) of CountBright^TM^ beads was added when cells had just been plated in a 96 wells plate after cell isolation,. During flow cytometry sample acquisition, the beads were counted along with cells (figure S3d and S4a). Because the total number of beads is known (given by the lot number), the number of cells per acquired sample (the absolute count) is obtained by relating the number of acquired cells to the number of acquired counting beads, then multiplying it by the number of total beads originally plated with the cells. At least 1000 bead events were acquired to assure a statistically significant sampling of sample volume. We then multiplied this absolute cell count per acquired sample, by the ratio this plated sample volume represented of the total cell suspension of the sample, e.g. if we plated one tenth of the sample we acquired with beads, we would multiply the absolute cell count/ acquired sample by ten to obtain the absolute cell count/total sample.

### Statistical inference of fluorescence loss probability

All HEK cells data in Fig. 2c and Fig. 2e for which the percentage of mScarlet-positive cells was less than 34.46% were excluded before processing. All data were fitted with a nonlinear mixed-effects model using ‘nlme’ package (vision version 3.1-163) in R (version 4.3.2) to estimate the probability of loss of m-Scarlet-expression per division^32^. The experiment number was treated as a random effect and the probability of losing m-Scarlet-expression was a fixed effect.

A bootstrap method was used to evaluate 95% confidence intervals^33^. In detail, a bootstrap sample was created by sampling with replacement from the experiment number and technical repeat of the original dataset and an estimate was obtained from the bootstrap sample via the model. This process was repeated 100,000 times to estimate the distribution of the estimates, from which basic upper and lower bootstrap confidence intervals were determined.

For Fig. 2d, Fig. 2f, and Fig. S2e-h ((Table S1), three-fold cross-validation was employed, where data from two of the three HEK293 experiments were used as training data while the data from the remaining experiment was used as testing data. Training data were fitted with a nonlinear mixed-effects model to estimate the probability of loss of m-Scarlet-expression. The experiment number was treated as a random effect and the probability of losing m-Scarlet-expression was a fixed effect.

The average division count was estimated for the testing data using the probability of loss of m-Scarlet-expression per-division from the training data. Linear regression was used to determine the relationship between the average cell population doubling and the average division count obtained by DivisionCounter using ‘stats’ (R version 4.3.2). A 95% confidence interval was computed using bootstrapping by sampling with experiment number and technical repeat from training data, and sampling with time from the testing data. The Pearson correlation coefficient was calculated using the ‘cor’ function with the method ‘pearson’ in R.

Fig. S2i-j and Fig. S3a plot the probability of loss of m-Scarlet-expression with division for MEF cells, MM468 cells and Nalm6 cells respectively. To estimate the probability of loss per-division, a nonlinear mixed-effected model was fit, and a 95% confidence interval was calculated using bootstrapping by sampling with experiment number and technical repeat. For fitting, all data for which the percentage of mScarlet-positive cells was less than 34.46% were excluded.

Fig. 2h displayed the average division count estimate calculated using the percentage of mScarlet-positive cells at 33°C in the DivisionCounter formula with the probability of losing the m-Scarlet-expression of the 37°C versus the population doubling computed from cell counts at 33°C. Linear regression was applied, and the 95% confidence interval was computed using bootstrapping by sampling with experiment number and technical repeat from 37°C data, and sampling with time from 33°C data. The Pearson correlation coefficient was calculated via the ‘cor’ function with the method ‘pearson’ in R.

In Fig. 2i, using the best estimate of the probability of losing the m-Scarlet-expression of HEK293T cells transfected with 26G-DivisionCounter cultured at 37°C as the parameter to fit the DivisionCounter formula, average division counts were predicted by the percentage of FP-positive cells. Simple linear regression and confidence intervals were evaluated in R. The output variable was average division count obtained with DivisionCounter, the input variable was time, and the intercept of this linear regression was 0.

The Spearman correlation coefficient in Fig. 2g was calculated by the ‘cor’ function with the method ‘spearman’ in R.

### DivisionCounter rationale

The DivisionCounter method consists of two parts: a lentiviral construct and a mathematical formula that, when used together, enable the estimation of the average division count of a cell population, i.e. the mean of the division counts of every cell in the cell population of interest. This development was influenced by (Weber et al., 2016), where it was mathematically established that the average division of a cell population could be inferred simply by tracking label proportion in a cell population that had a small likelihood of label loss at each division event. The only conditions were that cells should all be initially labelled, lose their label expression at each cell division with a fixed probability, and never reacquire it. Weber *et al.* therefore described a DivisionCounter formula for an irreversible two-state system (Weber et al., 2016).

However, our DivisionCounter formula describes the dynamics of a more involved three-state system, due to the use of microsatellite (MS) as the driver of label expression change. In search of a division linked mutative element to build our DivisionCounter, we decided to use MS due to their low mutation rate (between 10^2^ and 10^4^ per cell division) (Lai and Sun, 2003). MS mutations indeed generally occur by the addition or deletion of an entire nucleotide motif, which in our case is a single guanine. Therefore, by placing this MS between our DivisionCounter construct’s start codon and the mScarlet gene, MS mutation would cause the addition or deletion of one nucleotide, therefore altering the DNA sequence, and more importantly shifting our transgene mRNA translation reading frame by +1 or -1 before the mScarlet gene thus causing improper mScarlet translation leading to mScarlet expression stoppage. Hence, similarly to Weber *et al*., our DivisionCounter construct label expression would either be on or off, however the underlying mechanism of this label expression alteration comes from a three-state system, dictated by the biology of mRNA translation into protein depending on RNA sequences of nucleotides triplets, known as codons. The mathematical analysis in (Weber et al., 2016) relied on the irreversible label loss. Here, we derived a new three-state DivisionCounter formula that takes into account the biological specificity of reversibility in our experimental system.

### Microsatellite frameshift mutation probability: 3-states model

The first step in writing our DivisionCounter formula arises from analysis of a model of a three-state system that tracks changes of mRNA translation reading frame states and its link to division-induced MS frameshift mutation. We modelled our three-states system as a Markov chain that captures the reading frame state of a given cell through divisions amongst the three possible mRNA reading frames, namely in frame, +1 frameshift, -1 frameshift. All cells will be assumed to start in the “in frame” state and, at each cell division, will have a cumulative frameshift mutation probability, p, to move to +1 reading frame (after a guanine addition) or to move to a -1 reading frame (after a guanine deletion). If the cell is in state 1 and experiences a -1 mutation, it moves to state 3. Similarly, if a cell is in state 3 and experiences a +1 mutation, it moves to state 1. In this way, the state records the in- and out-of-frame state of the cell.

We defined q as the probability to move from the current state to the +1 reading frame state, and also to the -1 reading frame state, so that the probability of a mutation in either direction was p=2q. This means a cell will have a probability to remain in its current state equal to one minus to probabilities to undergo either frameshift mutations, which is equal to 1-2q. In mathematical terms, analysis of this system relies on the following Lemma.

### Lemma 1

Let

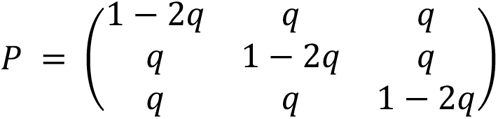

Then, for any n ≥ 1,

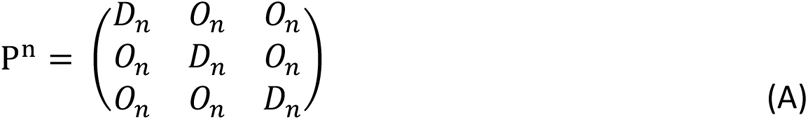

With 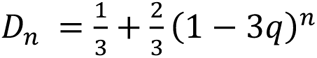 and 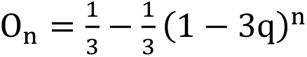.

**Proof**. Due to symmetries, P^n^ has the form in eq. (A) with D_1_ = 1 − 2q and 𝑂_1_ = 𝑞. As P^n^ = P ⋅ P^n-1^, we obtain the inductive relations:

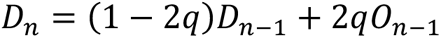

and

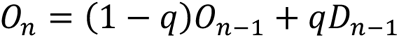

Subtracting the second equation from the first gives

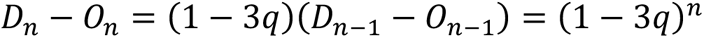

for all n ≥ 1. Hence, using one of the original inductive relations can be re-written as

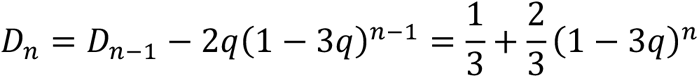

and the statement of the lemma follows.

As can be seen from the expressions for 𝐷_𝑛_ and 𝑂_𝑛_, the steady state equilibrium as is to have 1/3rd of cells be in the reading frame and 2/3rds of cells be out of the reading frame. We refer to this as the plateau.

Now that a three state reading frame system was modelled, we used it to derive an average division estimator formula, 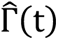, of a given cell population, Z(t), at time t, based on its percentage label expression, meaning the percentage of cells in the “in frame” state, Z_1 (t)/Z(t), where Z_1_(t) is the number of cells in “in frame” state at time t. In order to do so, the cell population Z(t) was defined as the sum of all cells in each reading frame state (1, 2 and 3) at time t, and was characterized by a vector of proportions of cells with different initial states, 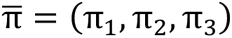, setting to π_1_ = 1 and π_2_ = π_3_ = 0 as all cells will start in “in frame” state as label positive cells. We will refer to the resulting formula as the General DivisionCounter formula (1).

We denote by 𝑍*_i_*(𝑡),the number of cells of type 𝑖 at time 𝑡, 𝑖 = 1,2,3, so that 𝑍(𝑡) = 𝑍_1_(𝑡) + 𝑍_2_(𝑡) + 𝑍_3_(𝑡) the total number of cells at time 𝑡. Then,

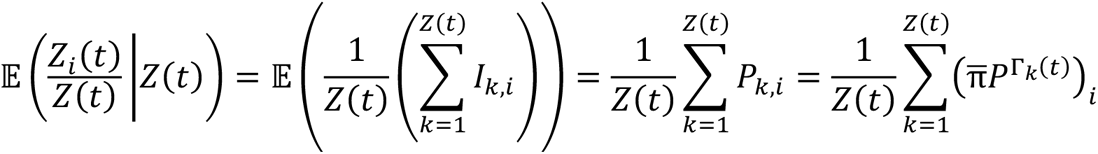

where 𝐼_𝑘,i_ is the indicator that cell 𝑘 is of type 𝑖 and 𝑃_𝑘,i_ is the probability that cell 𝑘 is of type 𝑖, and by 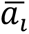 we denote the 𝑖-th element of a vector 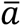 Then, taking expectations,

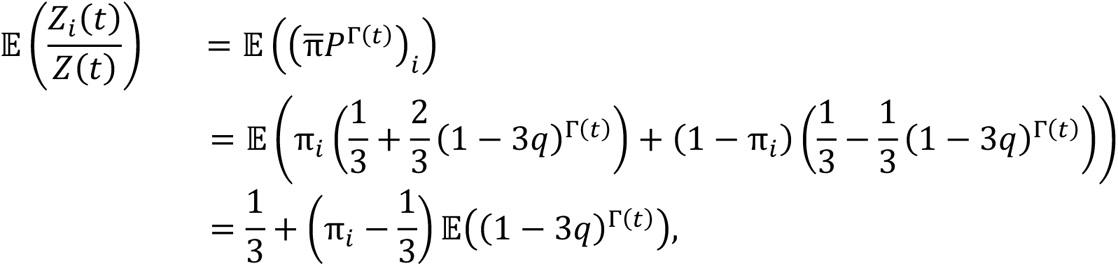

Or

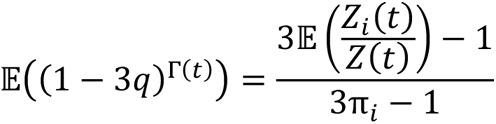

Hence, log 𝔼(𝑒^𝑠X^) = 𝑠𝔼(𝑋) + 𝑜(𝑠) for small *s*, one can use the estimator

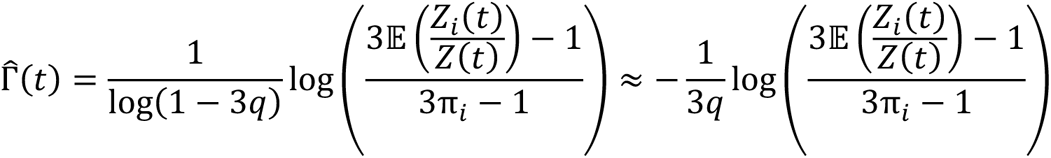

for *q* is smaller than 1/3. Note that, with π_1_ = 1 and 𝑞 = 𝑝/2, the estimator takes the form

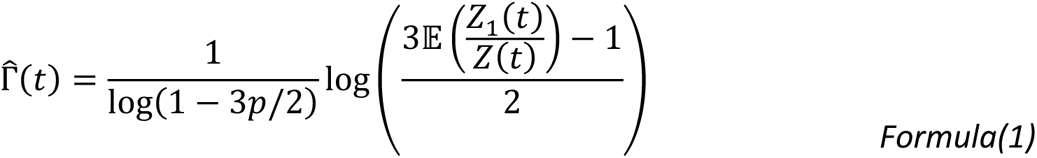

### Growth rate inference from cell numbers

Cell counts measured by flow cytometry with counting beads were the output data in Fig. 3f and Fig. 4c, and time points were the input data. The data were fitted by the exponential model using ’minpack.lm’ package (version 1.2-3) in R. Bootstrap samples were created by sampling with time to get 95% confidence intervals.

The day 20 BM and Spleen cell counts were predicted by day 7 and day 13 cell counts employing the best fit exponential model for BM and Spleen (Table S8). Permutation tests and Bonferroni correction were used to challenge the cell counts at day 20 and the predicted cell counts at day 20^34^. The null hypothesis for the permutation test for BM was that the mean number of cells at day 20 was independent of whether the cell counts were predicted or not. Monte Carlo approximation with 500,000 test statistics was used to estimate the p-value for the left-tailed test. The null hypothesis for the permutation test for Spleen was that the mean number of cells at day 20 was independent of whether the cell counts were predicted or not. Monte Carlo approximation with 500,000 test statistics was used to estimate the p-value for the right-tailed test.

Permutation tests were also used to observe differences in the Nalm6 cell count in BM of mice injected with CAR-T and Mock-T on the same day in Fig. 4c (Table S9). The null hypothesis was the mean of the Nalm6 cell count is independent of whether in different treatment on the same day. Monte Carlo approximation with 500,000 test statistics was applied to estimate the p-value for the left-tailed test, and Bonferroni correction was performed.

Permutation tests and Monte Carlo approximation were also used to statistically challenge differences in the cell count between different organs on the same day in Fig. 3c. The null hypothesis was the mean of the cell count is independent of whether in different organs on the same day. Table S2 reports the left-tailed p-value result, and Bonferroni correction was performed.

To challenge the null hypothesis that the fitted cell’s growth was independent of whether the cell in the 26G microsatellite or 26N microsatellite, a permutation test was used. A Monte Carlo approximation was used to estimate the left p-value. Table S3 showed the left-tailed p-value result, and Bonferroni correction was applied.

### Growth rate inference from DivisionCounter division counts

In Fig. 3d (Table S5) and Fig. 4e, using the probability of losing the m-Scarlet-expression expression of BFP+Lucipherase+Nalm6 cells infected with 26G-DivisionCounter *in vitro* as the parameter to fit the DivisionCounter formula, average division counts in vivo were predicted by the percentage of FP-positive cells. A simple linear regression was employed. The output variable was the average division counts obtained with DivisionCounter and the input variable was the time (days post-injection). A bootstrap sample with 100,000 datasets was created to get 95% confidence interval by sampling with experiment number and technical repeat from the *in vitro* dataset, and time from the in vivo dataset.

To challenge the hypothesis that division rates were independent of organs, permutation tests were employed (Table S6)^34^. Monte Carlo approximation with 500,000 test statistics was applied to estimate the p-value for the left-tailed test, and Bonferroni correction was performed.

Permutation tests and Bonferroni correction were used to statistically assess the average division counts obtained with DivisionCounter between different organs in Fig. 3e (Table S4). The null hypothesis was that on day 20 the mean of the average division count obtained with DivisionCounter is independent of the organ. The left-tailed p-values were calculated by Monte Carlo approximation with 500,000 sampling.

Permutation tests and Monte Carlo simulations were utilized to analyze the differences in Nalm6 average division counts obtained with DivisionCounter in BM of mice injected with either CAR-T or Mock-T cells on the same day, as illustrated in Fig. 4d. The null hypothesis was that the mean Nalm6 average division counts does not depend on the type of treatment, CART or Mock-T, on the same day. To evaluate the p-value for a left-tailed test, Monte Carlo simulations were performed with 500,000 iterations, followed by a Bonferroni correction for multiple comparisons.

### DivisionCounter in silico experiments

In silico simulation of cells exponentially dividing and changing their FP-expression changes over a set time were developed in R to investigate the properties and behaviour of the DivisionCounter.

At the start of each simulation, a cell population (n0) with all an “in frame” DivisionCounter construct are set to divide with exponential growth rate of ln(2)/D, where D is the doubling time (D). Here, D was set to one to mimic HEK293T cell growth. When cells divide, they have a frameshift mutation probability, p, of losing or gaining the label expression depending of its previous state. Label expression state of a cell is considered *off* if the DivisionCounter integrated construct reading frame states are “out of frame”, and *on* if the construct reading frame is in “in frame” state. Cells were then left to grow for 50 days during which the percentage of label positive cells in the cell population was recorded every half day. Cells were split to their starting cell count (n0) every 3 days. For the figures of the paper, we ran 100 simulations for the parameters detailed in Table S12.

## Data Availability

All datasets generated during this study are available at: https://github.com/changliu979/DivisionCounter and https://github.com/TeamPerie/Hustin-et-al

## Code Availability

All source code generated during this study is available at: https://github.com/changliu979/DivisionCounter and https://github.com/TeamPerie/Hustin-et-al

## Acknowledgments

We thank the Vallot team and the Menger team for providing us the MDA-MB-468 and Nalm6 cell lines. We thank all the staff from the Flow Cytometry Core Facility, in particular Lea Guyonnet and F. Tabarin and A. Batistella from the UMR168 Biologie Moléculaire et Biologie Cellulaire (BMBC) platform, Institut Curie, for their assistance. We thank Sheila Lopez-Cobo for helpful discussions for the Nalm6 experiment and Veronica A Raker for feedback on the manuscript. This work is part of a project that has received funding from the European Research Council (ERC) under the European Union’s Horizon 2020 research and innovation program 758170-Microbar and 101125069-Dynamyelo (to L.P.), and from the Fondation pour la recherche sur le cancer (ARC) (to L.H.). This publication has emanated from research conducted with the financial support of Science Foundation Ireland under grant number 18/CRT/6049 (to C.L and K.D). This project has received support under the program from the France 2030 launched by the French Government and from the Idex ANR-10-IDEX-0001-02PSL.

## Contributions

A.B. and L.H. designed the DivisionCounter constructs. A.B generated the plasmid constructs; F.T. produced the DivisionCounter lentiviruses with the help of L.H. L.H designed the *in vitro* experiments, with mentorship of L.P and comments from K.D, T.W, S.S. L.H and C.C performed the *in vitro* experiments. L.H, J.F, L.P designed the *in vivo* experiments with comments from S.M, C.C. L.H, J.F, and C.C performed the *in vivo* experiment. S.M prepared the CAR-T cells and performed *the in vitro* killing assay with supervision from S.A. L.H, L.P, C.C analyzed the flowjo data. L.H, C.L performed data analysis with supervision from L.P and K.D. C.L performed the statistical analysis with supervision from K.D. T.W, S.S and K.D developed the DivisionCounter formulae. L.H, C.L and S.S. wrote and performed the DivisionCounter simulations. L.P, L.H. and K.D wrote the manuscript with feedback from all authors.

## Extended information-Hustin et al

### Supplementary Figures

**Figure S1:**
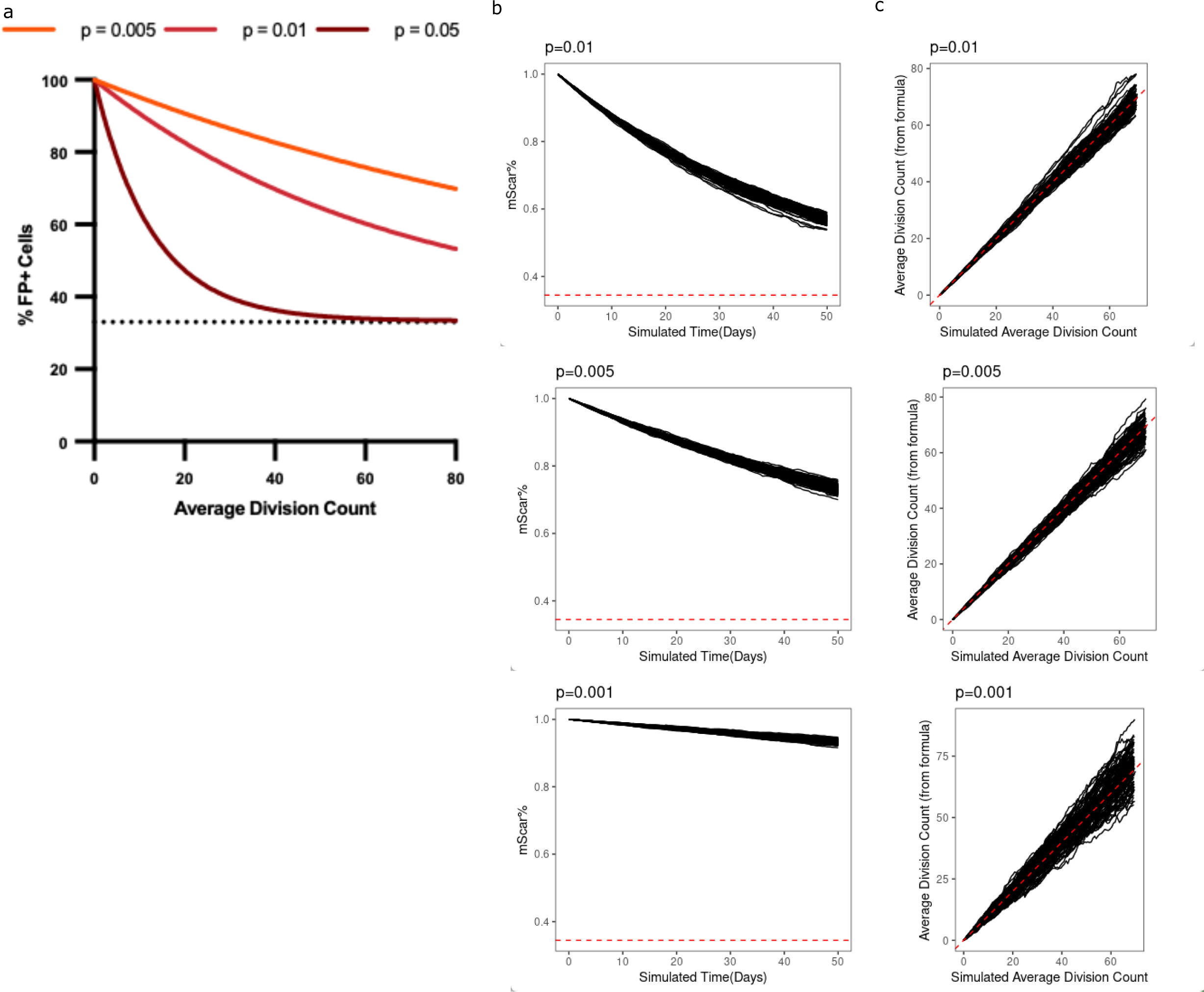
effect of the probability of losing FP expression on the DivisionCounter. a) Theoretical evolution of the percentage of FP+ cells as a function of division as defined by the DivisionCounter formula for different value of the probability of FP loss. b) Results of stochastic simulations of the percentage of FP+ cells over time for different probabilities of FP loss (p). N= 100 simulations. c) Average division count computed from the simulated % of FP+ cells against the true average division count in the simulations for different probabilities of FP loss. N= 100 simulations.

**Figure S2:**
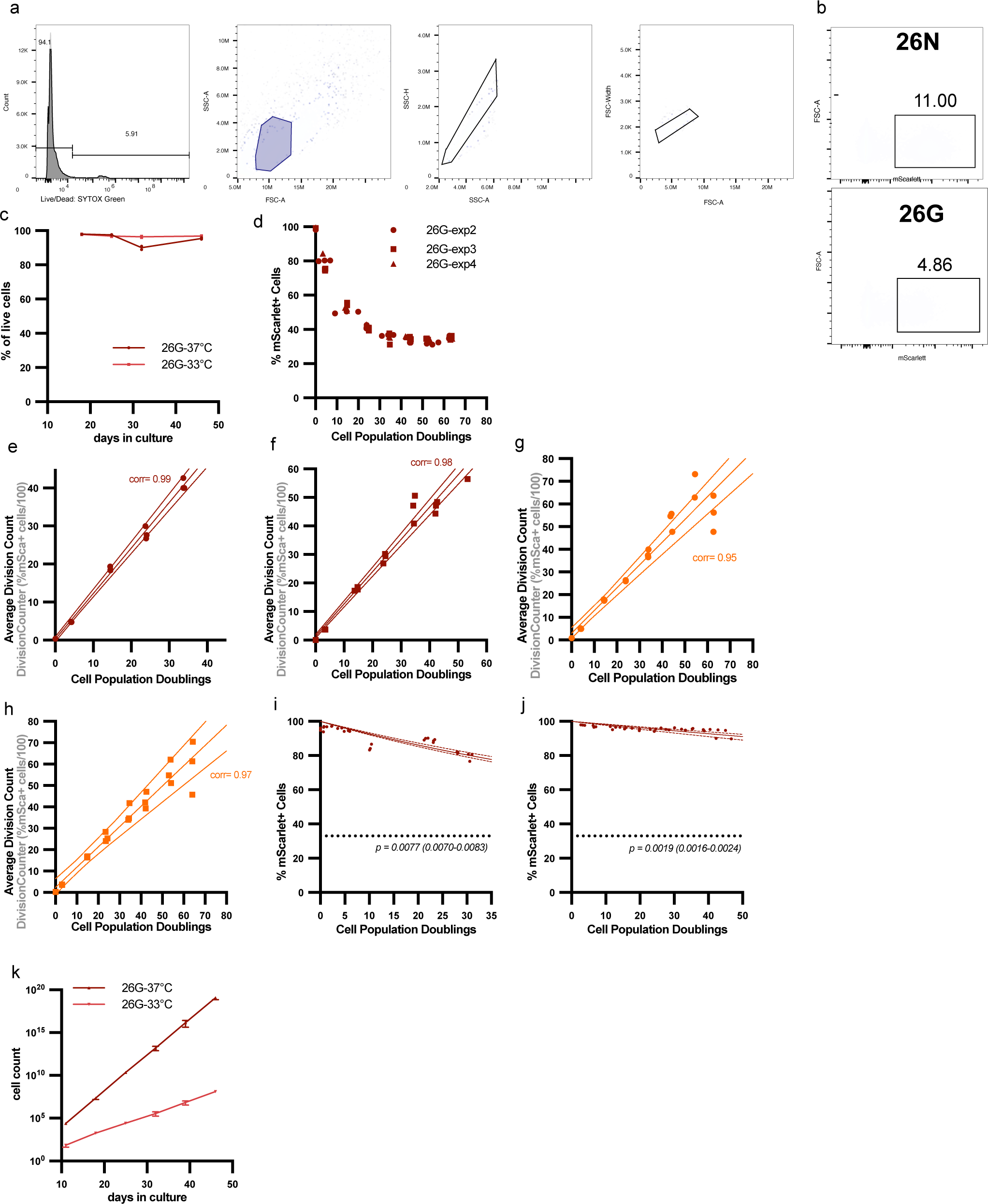
in vitro validation of the DivisionCounter method. a) First step of the gating strategy to sort and analyze DivisionCounter infected HEK293T cells. b) mScarlet gate used in the analysis and sort after the last gate of a) for the 26N- and the 26G-DivisionCounter. the percentage of mScarlet-positive cells indicated close to the box is used then to compute the division count using the DivisionCounter formula. c) percentage of HEK293T cells alive over time in culture at 37°C and 33°C. The live cells are defined as Sytox green negative cells as in the first panel of a) d) c) Percentage of mScarlet-positive cells over the number of divisions measured using cell counts (population doublings) for the 26G- DivisionCounter as in Figure 2c, colour-coded for different experiments. Each experiment (exp2, exp3, exp4) were done in triplicates. e) The computed average division count on the 2^rd^ experiment using the percentage of mScarlet-positive cells with the DivisionCounter formula and the probability fitted on the two other experiments for the 26G-DivisionCounter infected HEK293T (Table S1). The Pearson correlation is reported on the graph (corr). Each point represents independent triplicate acquisition. f) Same for the first experiment. g-h) Same than e-f for the 23G- DivisionCounter. i) Percentage of mScarlet-positive cells over the number of divisions measured using cell counts (population doublings) for the 26G-DivisionCounter in MEF cells sorted as 100% scarlet-positive at the start of the experiment. Each point represents independent acquisitions, from 2 experiments done in duplicates. The probability of losing mScarlet expression, p, was fitted using a linear model. The values shown are the best fit with confident interval obtained from bootstrapping. j) Same for MM468 cells. k) 26G-DivisionCounter infected HEK293T Cell counts over time in culture at 37°C (dark) or 33°C (light red) measured using the Septor cell counter. Mean and SD displayed from 3 experiments done in triplicates.

**Figure S3:**
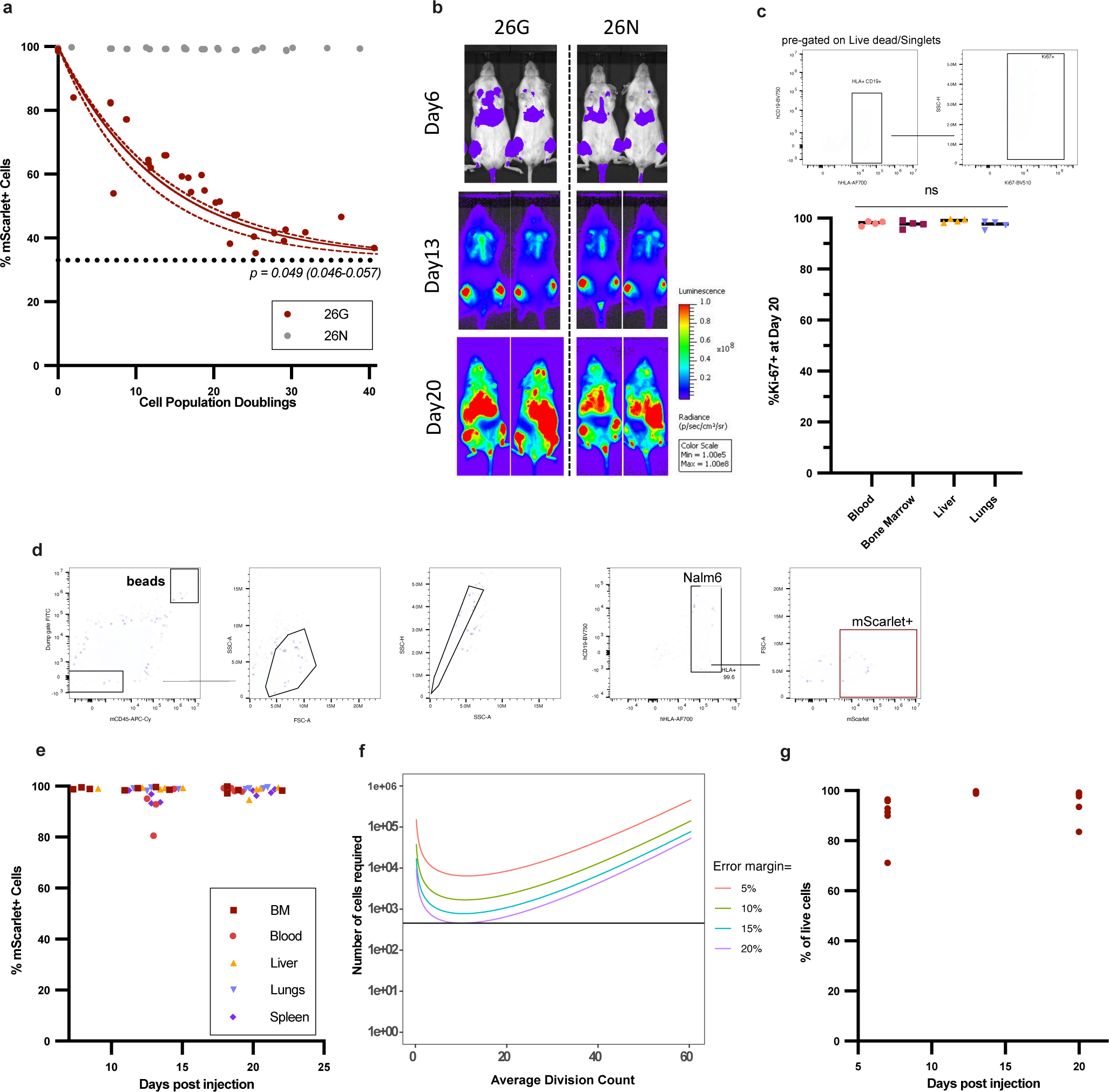
in vitro DivisionCounter Nalm6 cells experiment and in vivo bioluminescence, Ki67 and gating strategies. a) Percentage of Nalm6 mScarlet-positive cells over the number of divisions measured using cell counts (population doublings) for the 26G- DivisionCounter (red) and 26N- DivisionCounter (grey). For 26G-DivisionCounter, starting sorted cell population was either 100% scarlet-positive (dark red, mSca+) or 100% scarlet-negative (lighter red, mSca-). Each point represents independent acquisition from 3 experiments. The probability of losing mScarlet expression, p, was fitted using a nonlinear mixed-effects model. The values shown are the best fit with confident interval obtained from bootstrapping. b) representative pictures of total body bioluminescence measurements over time in the mice injected with 26G- and 26N-DivisionCounter infected Nalm6. 2 mice representative of 12 mice for day 6, 8 mice for day 13 and 4 mice at day 20. c) top: representative gating of Ki67 positive cells. lower: percentage of Ki67-positive cells at day 20 in the blood, BM, Liver and Lungs. Each point represents a different mouse, n=4 mice. d) representative gating strategy for the Nalm6 cells and mScarlet expression in Nalm6 in vivo. e) percentage of 26G-DivisionCounter mScarlet-positive Nalm6 cells over time in the BM (dark red), blood (light red), Liver (yellow), Lungs (lavender), Spleen (purple). Each point represents a different mouse, n=4 mice. f) the average division count versus the number of cells that must be acquired to ensure that the measured division count is within a given percentage of the true one. For p=0.049, if 455 or more cells are measured, there is at least a 97.5% chance of being with 20% of the true value. g) percentage of Nalm6 cells alive at day 6, 13 and 20 in the BM. The live cells are defined as Sytox green negative cells.

**Figure S4:**
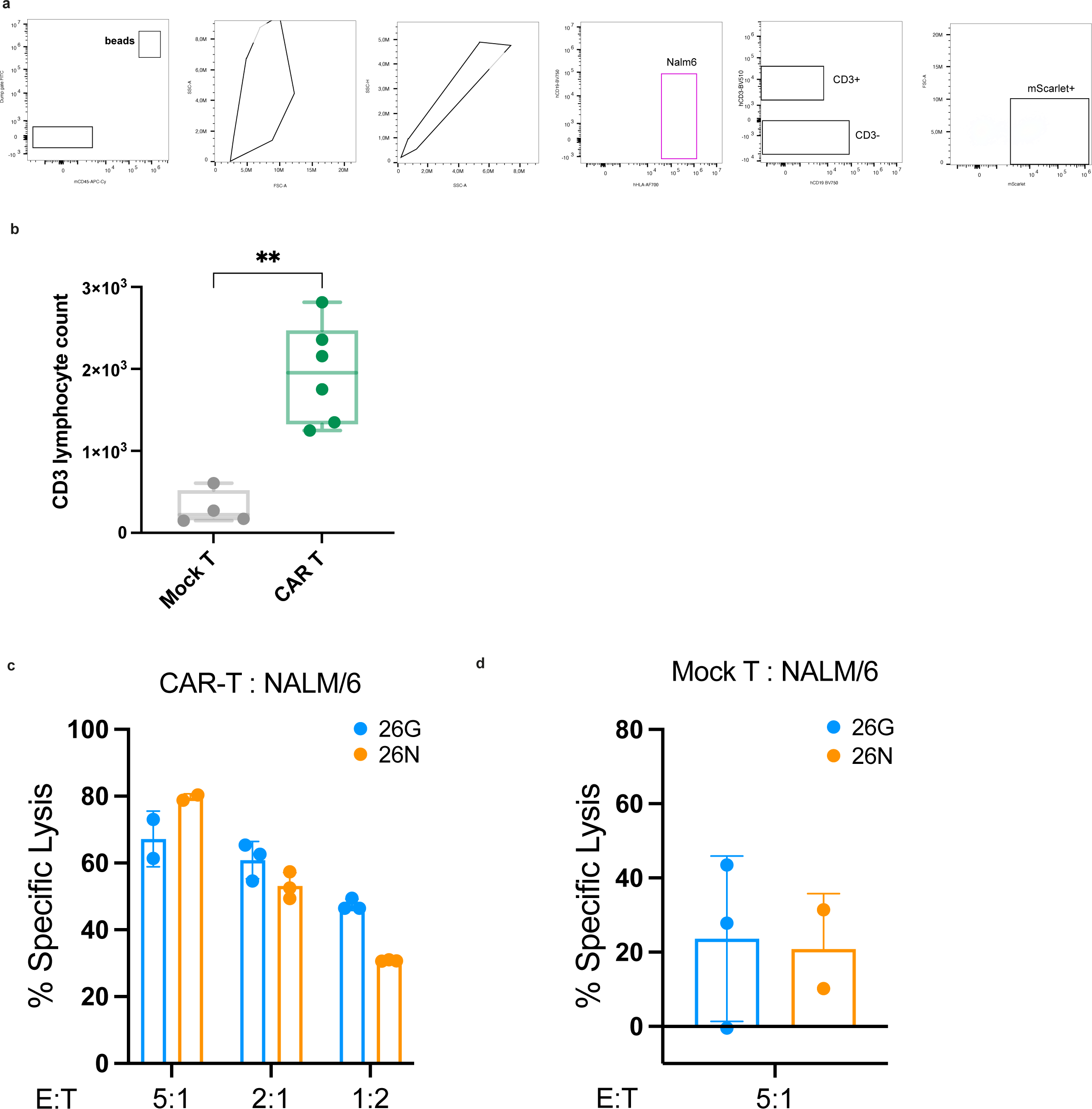
in vitro killing assay for CAR-T experiment and in vivo gating strategies. a) Luciferase-based cytotoxicity of 19-BBz CAR T cells in co-cultures with the 26G- or 26N-DivisionCounter infected Nalm6 cells at different ratios of T-cells (Effector, E) to Nalm6 cells (Target, T). Bars represent the mean. One experiment done in triplicates. b) same but with Mock-T. c) T-lymphoctes counts (CD3^+^) at day 7 post T-cell injection in the BM of mice injected with CAR-T (green) or Mock-T cells (grey). The cell number was scaled up to the bead ratio recovery and the volume of cells analysed. N=4 mice, ** indicates p<0.05 using an unpaired Mann-Whitney test. d) Representative gating strategy to define the Nalm6 cells, the percentage of mScarlet-positive Nalm6 cells and the CD3+ T lymphocytes.

### Supplementary Tables

**Table S1:** frameshift probability fitted on 2/3 experiments.

|  | fitted mutation probability |
| --- | --- |
| exp 3 & 4 | 0.048 (0.046-0.049) |
| exp 2 & 4 | 0.047 (0.044-0.049) |
| exp 2 & 3 | 0.046 (0.044-0.048) |

**Table S2:**
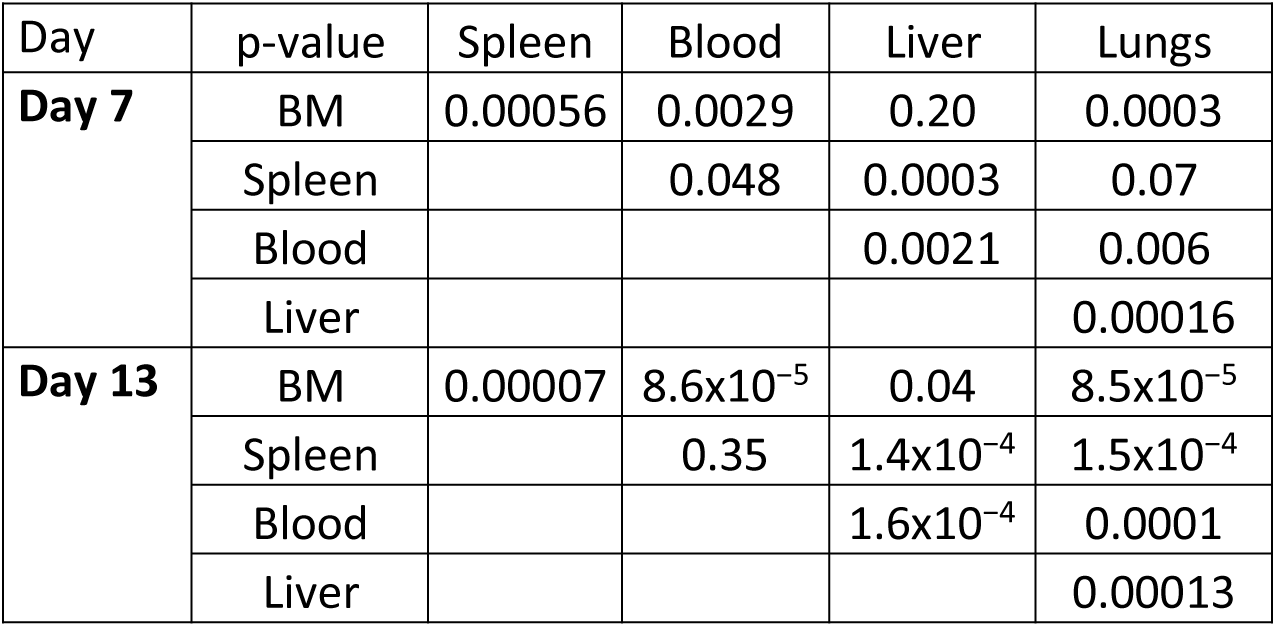
statistical comparison of cell counts across organs.

**Table S3:**
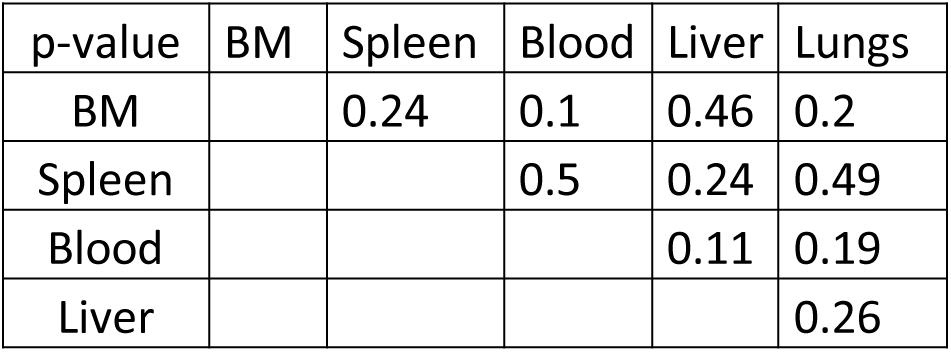
statistical comparison of fitted cell growth between 26G and 26N DivisionCounter.

**Table S4:**
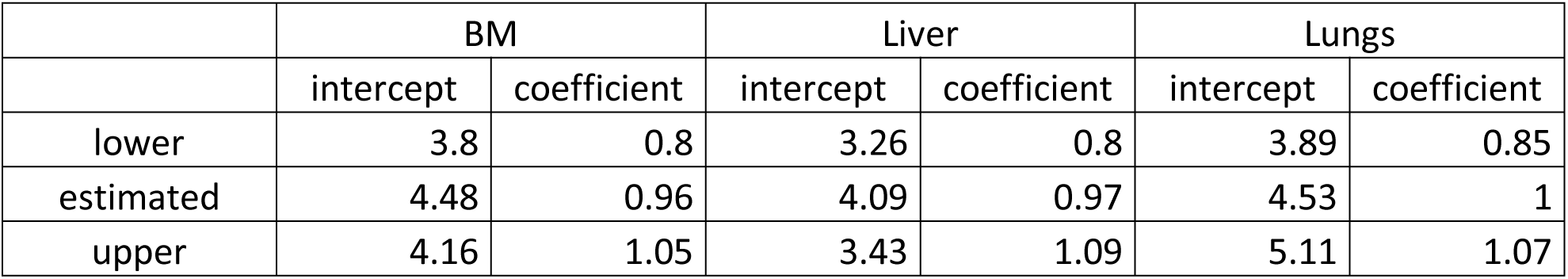
statistical comparison of the Nalm6 average division count between organs at day 20.

**Table S5:**
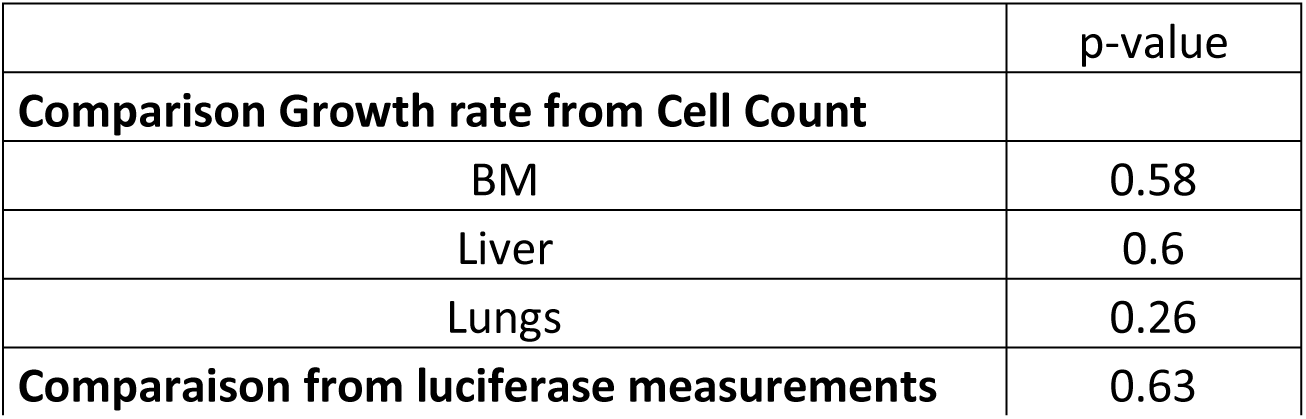
fitted Nalm6 division rate in each organ.

**Table S6:**
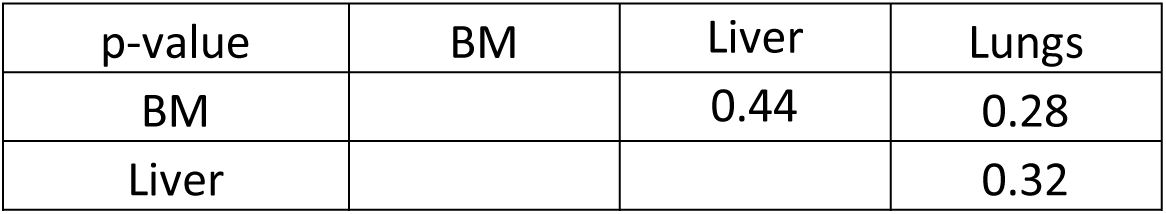
statistical comparison of the Nalm6 division rate between organs.

**Table S7:**
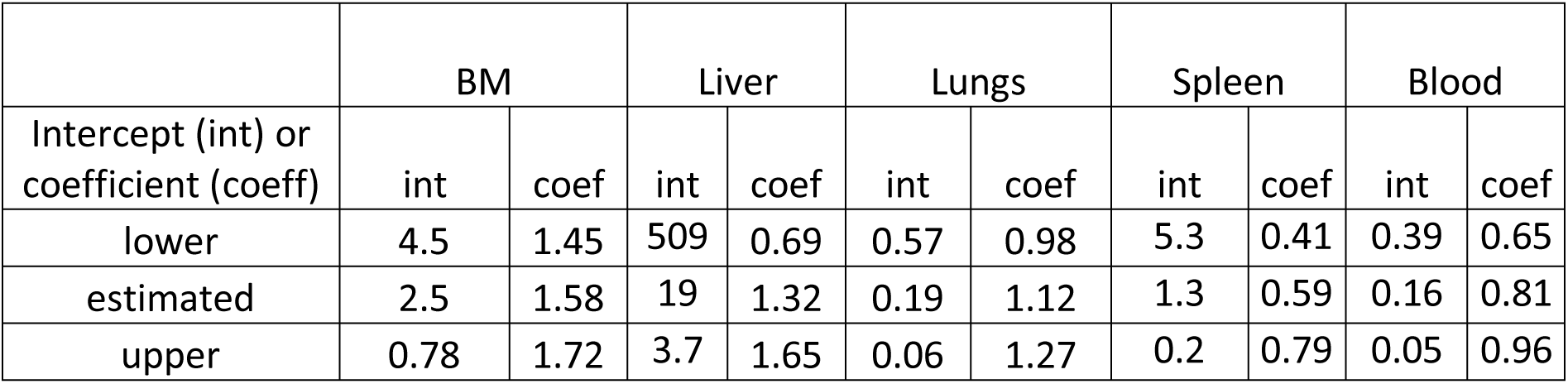
fitted net growth in each organ for day 7 and 13 based on cell counts.

**Table S8:**
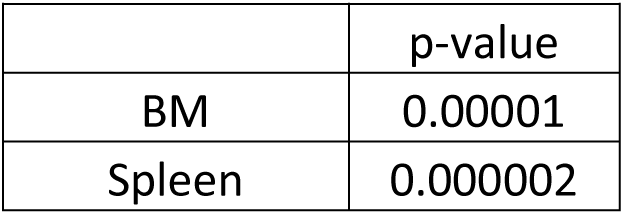
statistical comparison of the cell counts at day 20 for BM and Spleen compared to what is expected by exponential growth.

**Table S9:**
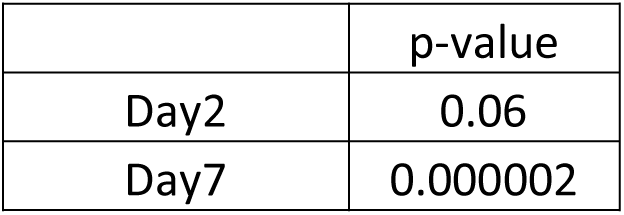
statistical comparison of cell counts between CAR-T and Mock-T treated mice.

**Table S10:**
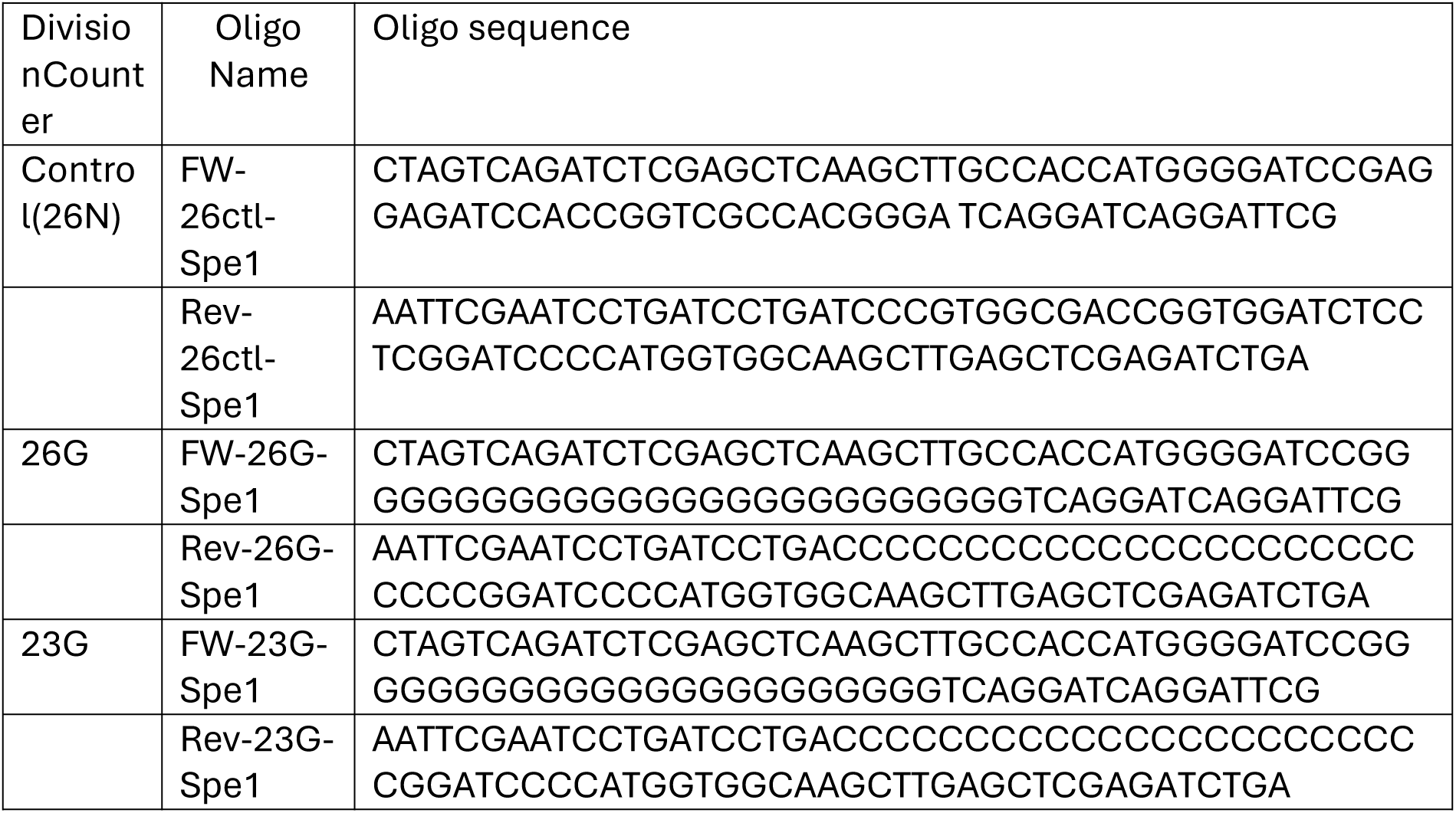
oligo sequence used for cloning.

**Table S11:**
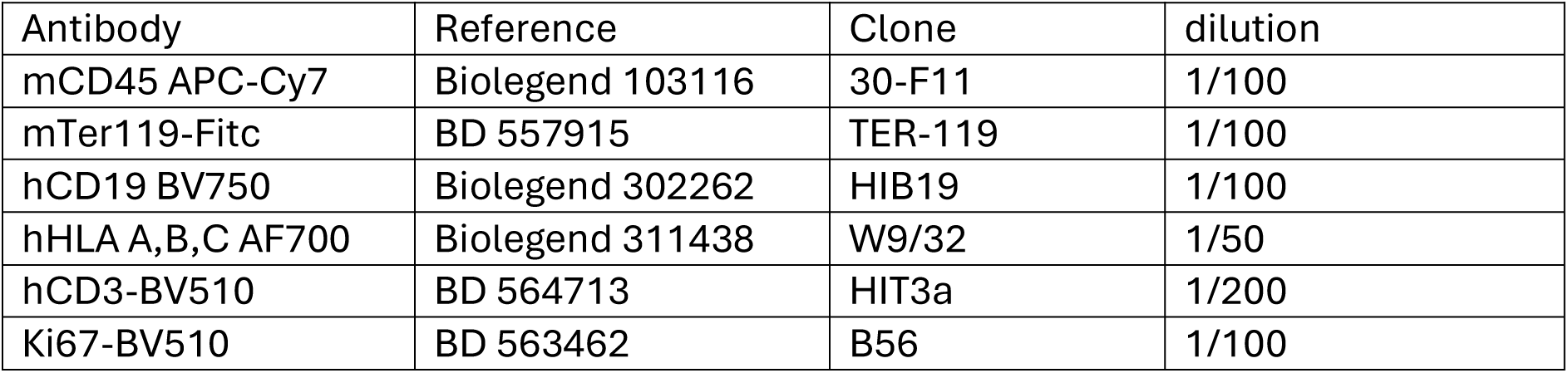
antibodies used for the in vivo experiment of figure 3 and 4.

**Table S12:**
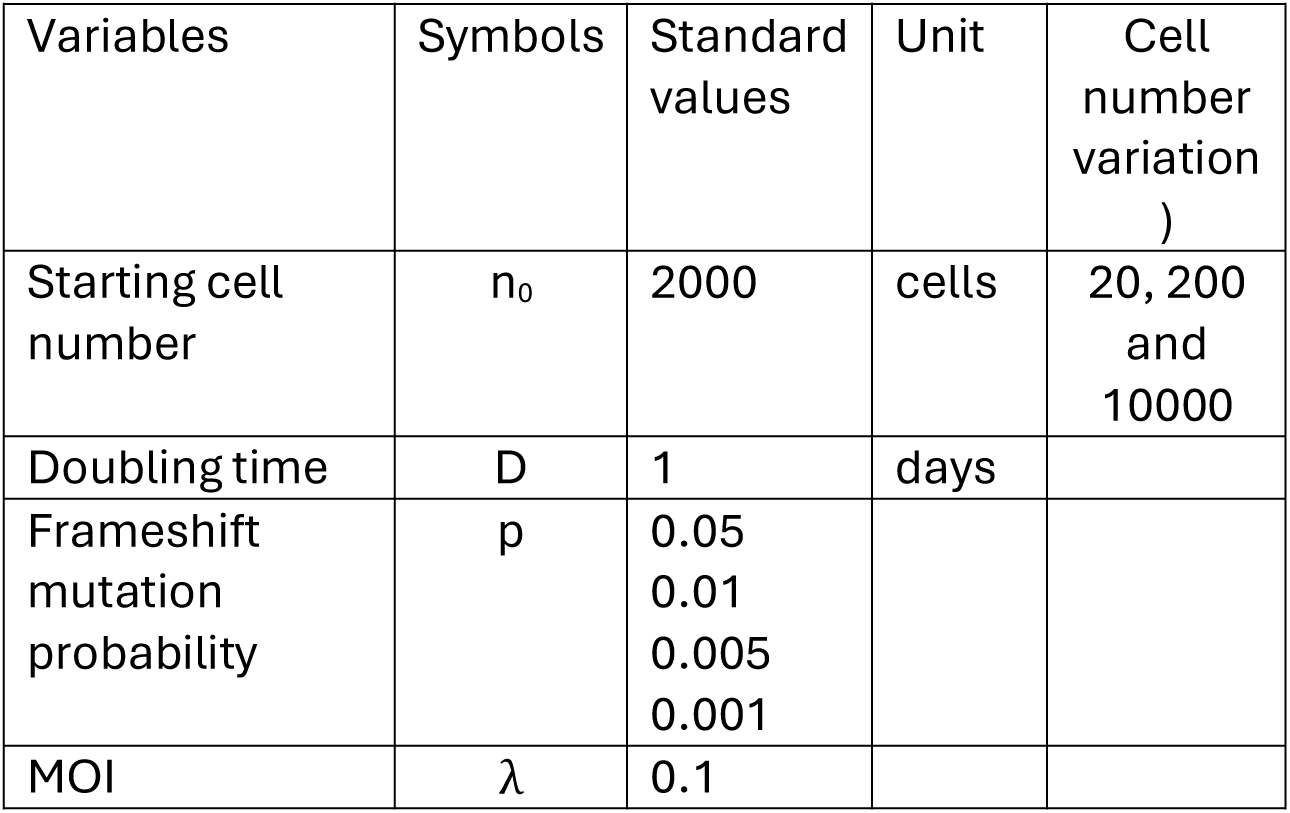
parameters used in the *in-silico* simulations.

